# Visual gamma stimulation causes prolonged enhancement of low-frequency blood flow oscillations across cortical regions in mice

**DOI:** 10.64898/2026.05.31.729102

**Authors:** Piergiulio R. Bressan, Emily Long, John Jiang, Rebecca Vithayathil, Zhixuan Guan, Yuwen Song, Bradley C. Rauscher, Nathan Chai, Kıvılcım Kılıç, Şefik Evren Erdener, Anna Devor, David A. Boas, Rockwell P. Tang

## Abstract

**Introduction:** Gamma entrainment using sensory stimuli (GENUS) uses 40Hz-pulsed sensory stimuli to entrain neural activity in the gamma band (30-150Hz). However, the effect of GENUS on low-frequency vascular oscillations has not been fully explored.

**Objectives:** The objective of this study is to elucidate the effect of GENUS on vasomotion in healthy mice and potential confounds for future application in disease studies.

**Methods:** Head-fixed, awake C57Bl/6 mice (n=18; 9M 9F) aged between 18 to 60 weeks were subjected to white light of either 40Hz visual flicker (GENUS), or constant stimulus (control). Blood flow was imaged using laser speckle contrast imaging (LSCI) before, during, immediately after 1 hour of stimulus, and 30min after the stimulus termination.

**Results:** A linear mixed-effects model showed that GENUS enhanced the magnitude of 0.2-0.4Hz blood flow oscillations by 38% during stimulation and by 30% at 30 minutes after stimulation compared to control when controlled for age, sex, and other factors. The effect on vasomotion was distributed across many cortical regions not limited to visual areas and lasted beyond 24 hours post-stimulus.

**Conclusion:** These results support the exploration of GENUS for increasing vasomotion in therapeutic contexts.

**Graphical Abstract:** 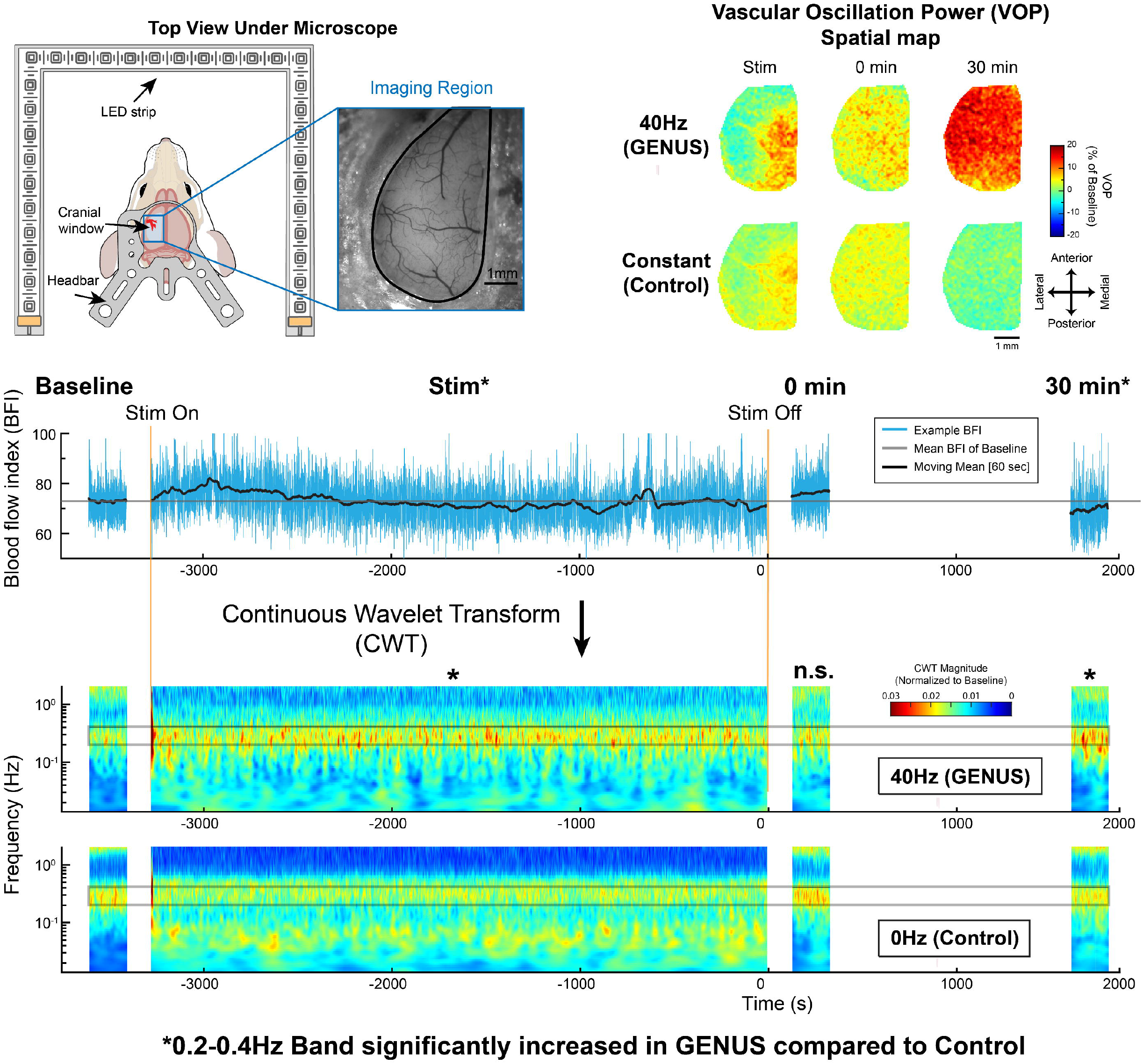

## Introduction

Low-frequency vascular oscillations (LFOs) have gained increasing interest in both physiological and pathological contexts [1-3]. These rhythmic variations in cerebral blood flow and hemodynamics are observed in a frequency band notably below cardiac and respiratory rates, but within the canonical frequency range (∼0.1Hz) of vasomotion [1,4]. During wakefulness, LFOs constitute a prominent component of spontaneous hemodynamic fluctuation in the cortex. Although it remains debated whether these slow vascular oscillations arise predominantly from intrinsic cerebral vasomotion or are partially driven by systemic arterial blood pressure fluctuations (Mayer waves) [5], they nonetheless influence cerebral blood flow dynamics and resting-state functional connectivity [6]. Arterial LFOs have additionally been show to facilitate perivascular clearance: in slow wave sleep, increased norepinephrine-driven LFOs drive glymphatic flow and waste clearance [7,8].

Glymphatic dysfunctions and perfusion deficits observed in Alzheimer’s disease [9-12] and stroke [13-17] have been linked to neural impairments, highlighting the potential of LFOs as a target for therapies for improved perfusion or waste clearance. Dysfunction of LFOs after acute ischemic injury may contribute to impairments in flow regulation and perivascular clearance in the early post-ischemic brain [13], processes which contribute to edema formation and the clearance of metabolic byproducts that can be damaging to spontaneous recovery [16]. Experiments in ischemia-reperfused mice have shown that, in the subacute phase (days to weeks after ischemia), LFOs are increased and positively correlated with long-term behavioral outcome [18], possibly facilitating recover via improved waste clearance. Supporting this link, glymphatic flow has been shown to facilitate post-stroke structural and functional recovery [13-16]. These findings motivate therapeutic interventions that support and enhance low-frequency vascular dynamics, offering a pathway to improved post-ischemic recovery.

Cerebrovascular dynamics at frequencies below 1 Hz can be cardiac, respiratory, myogenic, neurogenic or endothelial in origin [19]. While LFOs fall between the myogenic and respiratory bands, recent studies have linked them to neural activity through neurovascular coupling and neuromodulatory activity [20-22]. Many neurotransmitters and neuropeptides are vasoactive with receptors on arteriolar smooth muscle cells [23] where the majority of ∼0.1Hz LFOs originate [24]. In the cortex of awake mice, spontaneous LFOs were shown to correlate with and could be driven by fluctuations in the gamma-band power of neural electrical activity [20]. This raises the possibility that interventions that directly or indirectly entrain the gamma-rhythm could modulate LFO power and synchrony, and therefore perfusion and perivascular transport processes relevant to disease.

One such approach is Gamma ENtrainment Using Sensory stimulation (GENUS) [25,26], a non-invasive method that commonly relies on audiovisual stimuli modulated at 40 Hz to drive gamma-band neural activity. Research in both animals and humans has demonstrated that the resulting entrainment is not limited to primary sensory cortices, but extends to other brain regions such as the hippocampus and prefrontal cortex [25,27,28]. GENUS has already shown significant potential in ameliorating an underlying pathology in a mouse model ofAlzheimer’s disease by promoting Aβ clearance through enhanced glymphatic flow, with increases in both CSF influx and ISF efflux in the cortex [25,26,29]. However, the physiological mechanism linking gamma-band stimulation to enhanced glymphatic transport remains incompletely understood. Given that LFOs have been correlated with both gamma-band fluctuations [20] and clearance [7,8,30], these vascular oscillations might represent a missing mechanistic link between GENUS and its effects on glymphatics. In this study, we therefore hypothesize that gamma-band visual stimulation increases the power of LFOs in the cortex of awake mice. As application of gamma-band sensory stimulation in disease models other than Alzheimer’s, such as ischemic stroke, is still limited, gaining insight into a vascular effect can encourage its investigation in other disease models and thereby improve its clinical potential.

To evaluate whether visual GENUS can alter cerebrovascular dynamics in a way relevant to post-ischemic recovery, in this study we explore the effects of gamma-band visual stimulation on cortical LFOs in awake mice of varying age and sex. Throughout this study, we refer to LFOs as 0.2–0.4 Hz fluctuations in cortical blood flow, in accordance with both previous literature [8,18,24,31] and empirical observations in our data. Specifically, we quantify changes in 0.2–0.4 Hz LFO magnitude across cortical areas and pre- and post-stimulation timepoints, comparing the effects of rhythmic 40 Hz flickering white light stimulation to an equiluminant control light. GENUS stimulation robustly increased LFOs relative to pre-stimulus baseline as well as the control group, with the effect lasting up to 24 hours post-stimulus and spanning all regions which were imaged. By establishing the modulation of LFOs by GENUS, this study tests a candidate physiological mechanism that may link gamma-band sensory stimulation to enhanced perivascular transport, and motivates future investigation of GENUS in ischemic stroke models.

## Materials and Methods

### Experimental Design

#### Animal Preparation

All experiments and animal procedures were approved by the Boston University Institutional Animal Care and Use Committee and conducted in accordance with the Guide for the Care and Use of Laboratory Animals. A total of n=18 wild-type mice (9 male, 9 female; Jackson Labs C57BL/6J) between the ages of 18-60 weeks were used in this study. Group sizes were estimated by Lamorte power calculation (Supplementary Table ST1). Consistent with prior studies [18,32], mice underwent craniotomy and surgical implantation of a glass cranial window (Warner Instruments coverslip; approx. 3.5 x 6.5mm) over one cortical hemisphere at 14-30 weeks of age to enable optical imaging. A metal headbar was affixed to the cranium for head fixation. Following 2 weeks of surgical recovery, mice were habituated to head fixation by head-fixing daily for gradually increasing durations until each animal could remain head-fixed without signs of distress such as vocalizations for 120 minutes. Imaging was conducted between 2-37 weeks after craniotomy. Mice were inspected prior to each experiment for eye occlusions such as cataracts, in which case they would be excluded from further experimentation. However, no mice were excluded. Mice were housed in normal light cycles with ad libitum access to food and water. We used the ARRIVE checklist when writing our report [33].

#### In vivo imaging

Awake mice were head-fixed via the implanted headbar onto a custom cradle (Thorlabs) under a microscope setup as previously reported [34], surrounded by the stimulation setup (Figure 1A). Baseline vascular dynamics were recorded for 10 minutes using laser speckle contrast imaging (LSCI). Mice were then exposed to either 40 Hz flickering light (GENUS) or 0Hz constant light (control) for 1 hour, during which LSCI data were continuously acquired. 10-minute recordings of LSCI were acquired again at 0 minutes and 30 minutes after the end of stimulation. Mice were returned to their home cages after these measurements. During each LSCI measurement, mouse behavior was simultaneously recorded via an infrared camera. GENUS and control stimulation were performed repeatedly on mice, with a period of at least 48 hours between to prevent carryover effects. A subset of mice underwent an additional LSCI recording at 24 hours after stimulation to assess prolonged effects. A total of n=18 GENUS and n=32 control datasets were gathered, which included n=11 GENUS datasets and n=24 control datasets that had data at 24h. In 6 mice (2M, 4F), additional measurements were taken at 48, 72, 96, and 120h after control stimulation to track longitudinal variability. Group allocation was not controlled or anonymized during experiments and post-hoc analysis. Measurements were taken during the light phase, but the time of day was not controlled.

**Figure 1.**
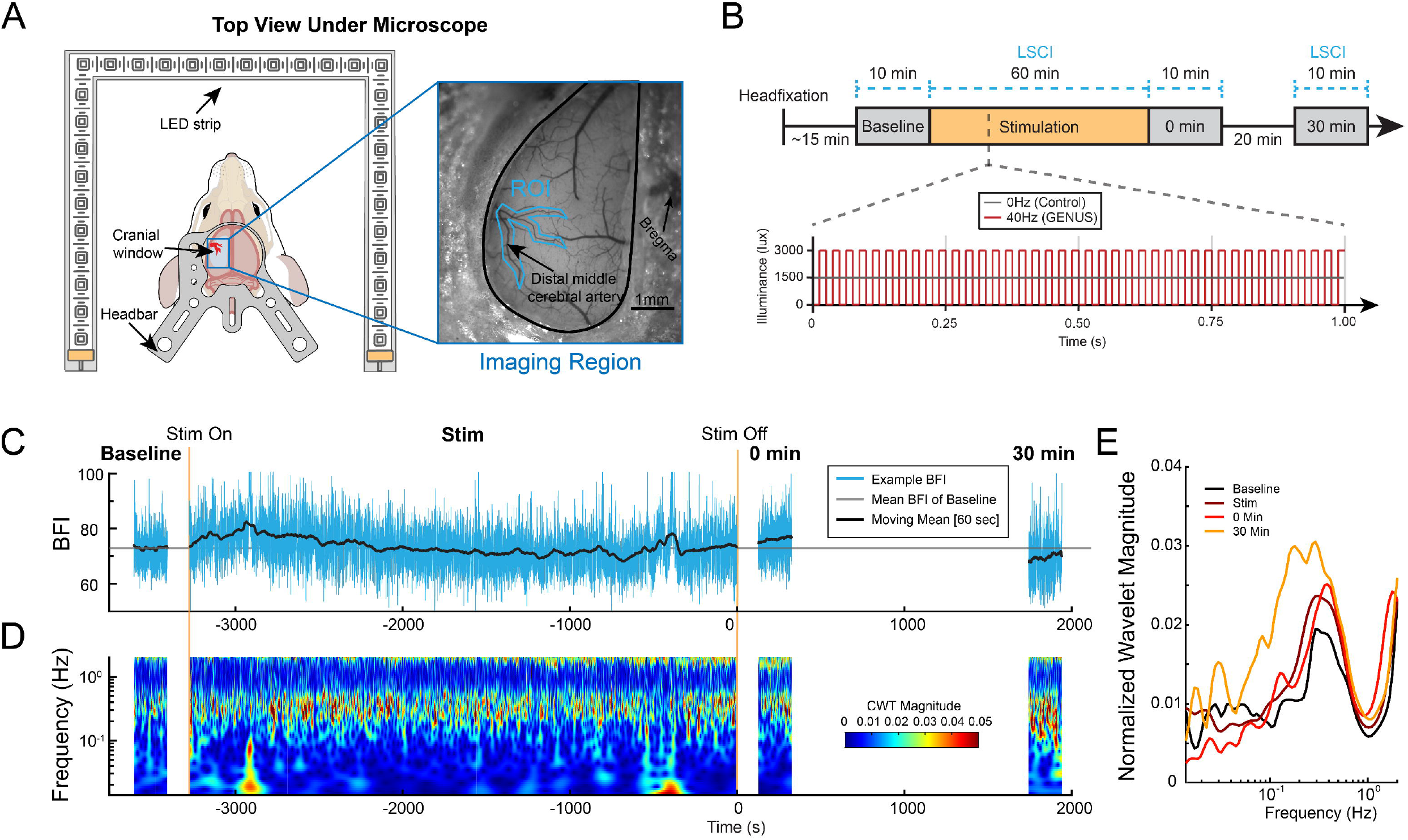
Blood flow dynamics were recorded before, during and after visual stimulus. (A) Diagram of the setup. Mice implanted with a chronic cranial window are headfixed for laser speckle contrast imaging (LSCI). The mouse is surrounded on three sides by an eye-level LED strip which is controlled during stimulation. During analysis, an arterial region of interest (ROI, teal) is selected for quantification of blood flow index (BFI). (B) Timeline of the measurement. One hour of visual stimulus (either 40Hz GENUS or 0Hz control) is delivered to the head-fixed mouse. LSCI is recorded in periods Baseline, Stim (Stimulation), 0min and 30min. (C) Example time series of BFI in the selected arterial ROI, with the trend marked in black for visibility. (D) Continuous wavelet transform (CWT) magnitude of the same BFI timeseries, showing changes in magnitude at different frequencies of the blood flow. (E) The CWT spectra of the BFI of each time period for the example data.

#### Stimulation setup

A custom visual stimulus setup was used. An LED strip (Pautix) was switched at either 40Hz (50% duty) or 0Hz by an Arduino Uno with a custom program and a separately powered MOSFET driver. Light intensity was controlled by pulse-width modulation at 490Hz and adjusted to a safe maximum exposure of 3000 lux. GENUS stimulation flickers between 0 and 3000 lux, while control stimulation is adjusted to 1500 lux to deliver similar total amounts of light. During stimulation, the LED strip is mounted approximately 6 inches from the mouse at eye level and surrounds the mouse on the front and sides. Stimulation was performed in a dark room for 1 hour per session.

#### Laser speckle contrast imaging (LSCI)

A custom LSCI system was used to measure cerebral blood flow (CBF) across the cortical window. The cortical surface was illuminated using a 785 nm VHG-stabilized laser (LP785-SAV50; Thorlabs) at a power density of 10 mW/cm^2^ and imaged by a CMOS camera (Basler acA2040-90μm NIR) through a 2× objective (TL2X-SAP; Thorlabs). A dichroic mirror, polarizer, and iris were placed in the optical path to optimize image quality. For the study of the vascular effects of GENUS stimulation, LSCI data was acquired at a framerate of 10 Hz at 5ms exposure for 10 minutes at each of the Baseline, 0min, 30min and 24h timepoints. Data was also acquired at 10Hz during the 1-hour stimulation period, except for 7 of the GENUS and 7 of the Control datasets where the framerate was reduced to 2 Hz during the hour-long stimulation period to reduce data volume. These datasets were interpolated to 10Hz before subsequent analysis. For the analysis of isoflurane effects on vascular oscillations, LSCI data was collected at a framerate of 10 Hz for 1 hour. During each measurement, an infrared behavioral camera (Svpro OV2710) was positioned laterally to the right eye for simultaneous whisker recording at 30 frames per second, using the LSCI illumination for contrast.

### Data and Statistical Analyses

#### Data processing

All data were analyzed using custom scripts written in MATLAB (MathWorks) and Rstudio. Raw speckle images were converted to speckle contrast (*K*) maps by calculating the ratio of the standard deviation to the mean intensity across a 7×7 pixel window. These were then used to compute blood flow index (BFI) with BFI = 1 / *K*^2^. Regions of interest (ROIs) were manually selected based on visible cortical vasculature beneath the cranial window. ROIs typically encompassed distal branches of the middle cerebral artery (MCA) along with adjacent parenchymal tissue. Identical ROIs were selected for each timepoint within a dataset. For each 10-minute LSCI recording, 200 seconds of resting state data with minimal motion were selected for further analysis. This selection aimed to eliminate potential confounds introduced by spontaneous movement or behavioral agitation. From the selected segments, a time series of the mean blood flow over the ROI was extracted.

To assess vascular oscillations, a continuous wavelet transform (CWT) was applied to each BFI time series (MATLAB cwt; 10 voices per octave, analytical Morlet wavelet, L1 normalization) after normalizing it by its mean. The mean magnitude of the transform was quantified within the 0.2-0.4 Hz frequency band, corresponding to low-frequency oscillations (LFOs), resulting in a LFO timeseries. For further statistical analysis, the mean of this time series was recorded as the scalar VOP of each recording period.

To assess mouse whisking, the motion energy over the right whisker pad was calculated by taking the absolute difference between adjacent frames before averaging over a manually selected ROI. Frames where the motion energy rose above 1 standard deviation from the 10-second moving mean were classified as whisking. Finally, the fraction of frames from the 200s window in which the mouse was whisking was calculated.

#### Spatial maps

Spatial maps of BFI and VOP were calculated for each timepoint of each dataset. For each pixel, the BFI time series was averaged over the same time range selected in the initial analysis. For each pixel, a wavelet magnitude value was calculated similarly to above, forming a map of 0.2-0.4Hz VOP. These maps were manually registered to the Allen Atlas based on lambda and bregma markings visible in widefield images, using custom MATLAB scripts. Each map for each timepoint was registered to a single reference image for each mouse, and the pixelwise percent change from baseline was calculated. All maps were masked to contain only pixels under the cranial window. After averaging registered maps for each treatment group and timepoint, only the region of pixels with maximum overlap between mice was kept.

A parcellation was performed using the Allen Atlas [35] by averaging BFI or VOP for each classified brain region. Brain regions with less than 20% area of overlap with the cranial window were discarded from further analysis. Regions that were present in less than 80% of the datasets were also discarded. Finally, the group mean BFI and VOP were calculated for each brain region at each timepoint and treatment group.

#### Linear mixed-effects modeling

Due to the possibility that BFI and VOP could be influenced by subject factors such as age and sex, linear mixed-effects (LME) models were used for rigorous post-hoc statistical analysis. First, the scalar BFI or VOP data were log-transformed to improve their normality, as the original data failed the Shapiro-Wilk normality test (Supplementary Figure S1, S2). The response variables were computed as 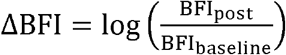 and 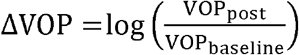, where BFI_post_ and VOP_post_ are the data from each post-stimulation timepoint, and BFI_baseline_ and VOP_baseline_ are the data from the baseline period. For all models, factors including treatment type, timepoint, the number of previous sessions for that mouse, age, number of days after craniotomy, and sex were considered. A random intercept (mouse ID) model was chosen, as in every case a random slope model had a singular fit without an increase in performance. The full model initially considered was:

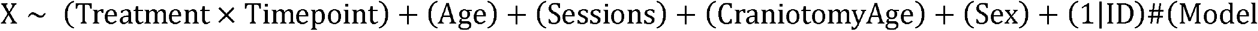

where X was ΔBFI or ΔVOP.

Age, session number, and days post-craniotomy were collinear (Supplementary Table ST1). To reduce model complexity, only one or less of these variables was included in each model. Three variations of Model 0 with only one of the three age variables were compared, to select the one which had the most significant effect. For the model of ΔBFI, none of the variables were significant (Supplementary Table ST2), so all three were omitted from the final model:

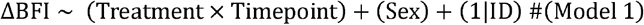

The random and fixed effects of Model 1 are provided in Supplementary Table ST3. Normality and homoscedasticity of the residuals is shown in Supplementary Figure S3.

For the first model of ΔVOP, all age variables were significant. As craniotomy age had the most significant influence on model performance (Supplementary Table ST4), session number and age were removed as variables from the final model:

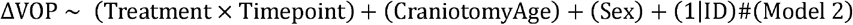

The random and fixed effects of Model 2 are provided in Supplementary Table ST5. Normality and homoscedasticity of the residuals is shown in Supplementary Figure S4.

For the analysis of ΔVOP at 24 hours post-stimulation, a separate model was used because only a subset of datasets included these data. Because other variables were not significant, the final model was:

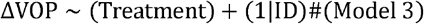

A t-test on estimated marginal means from each LME model (referred to herein as LME model t-test), with Satterthwaite approximation for degrees of freedom and Tukey adjustment for multiple comparisons, was used to assess significance of effects. The t-tests performed on Model 1, 2, and 3 are summarized in Supplementary Tables ST6, ST7, and ST8 respectively. Confidence intervals were estimated by *CI* = *<u>x</u>* ± *t*_*c*_ × *SE*, where *t*_*c*_ is the critical value of the t-distribution and *SE* is standard error. The code for LME modeling was implemented in R.

## Results

Awake, head-fixed mice (n=18 mice) were subjected to 1 hour of 40Hz (GENUS; n=18 datasets) or 0Hz (control; n=32 datasets) white light stimulation using a custom microscope and LED strip setup (Figure 1A). Laser speckle contrast imaging (LSCI) at 10Hz framerate and 5ms exposure was performed immediately before, during, and 0 and 30 minutes after stimulation (named Baseline, Stim, 0min, 30min respectively) (Figure 1B). Blood flow index (BFI) image series were calculated using spatial speckle contrast and averaged over a region of interest (ROI) encompassing distal branches of the middle cerebral artery (MCA) (Figure 1A) to get a BFI time series for each dataset (Figure 1C). The time series was analyzed using a continuous wavelet transform (CWT) (Figure 1D), which showed a peak in the wavelet magnitude around ∼0.3Hz (Figure 1E). Therefore, the 0.2-0.4Hz frequency band was selected for further analysis.

A similar analysis to Figure 1 was conducted for each dataset acquired. To allow for comparison between datasets, each BFI time series was normalized by subtracting the mean BFI of its Baseline. The spaghetti plot of the percent change in BFI shows that blood flow remained within ±10% of Baseline mean for most datasets (Figure 2A). When averaged for each recording period, there was a slight negative trend and slight positive trend in Stim and 0min, respectively (Figure 2B). However, there were no significant differences in BFI between treatment groups or at each time period compared to Baseline (LME model t-test). Significance levels were determined from t-tests on the estimated marginal means derived from a linear mixed-effects (LME) model that accounts for animal-specific variability and experimental covariates (see Methods and Figure 3 for model details).

**Figure 2.**
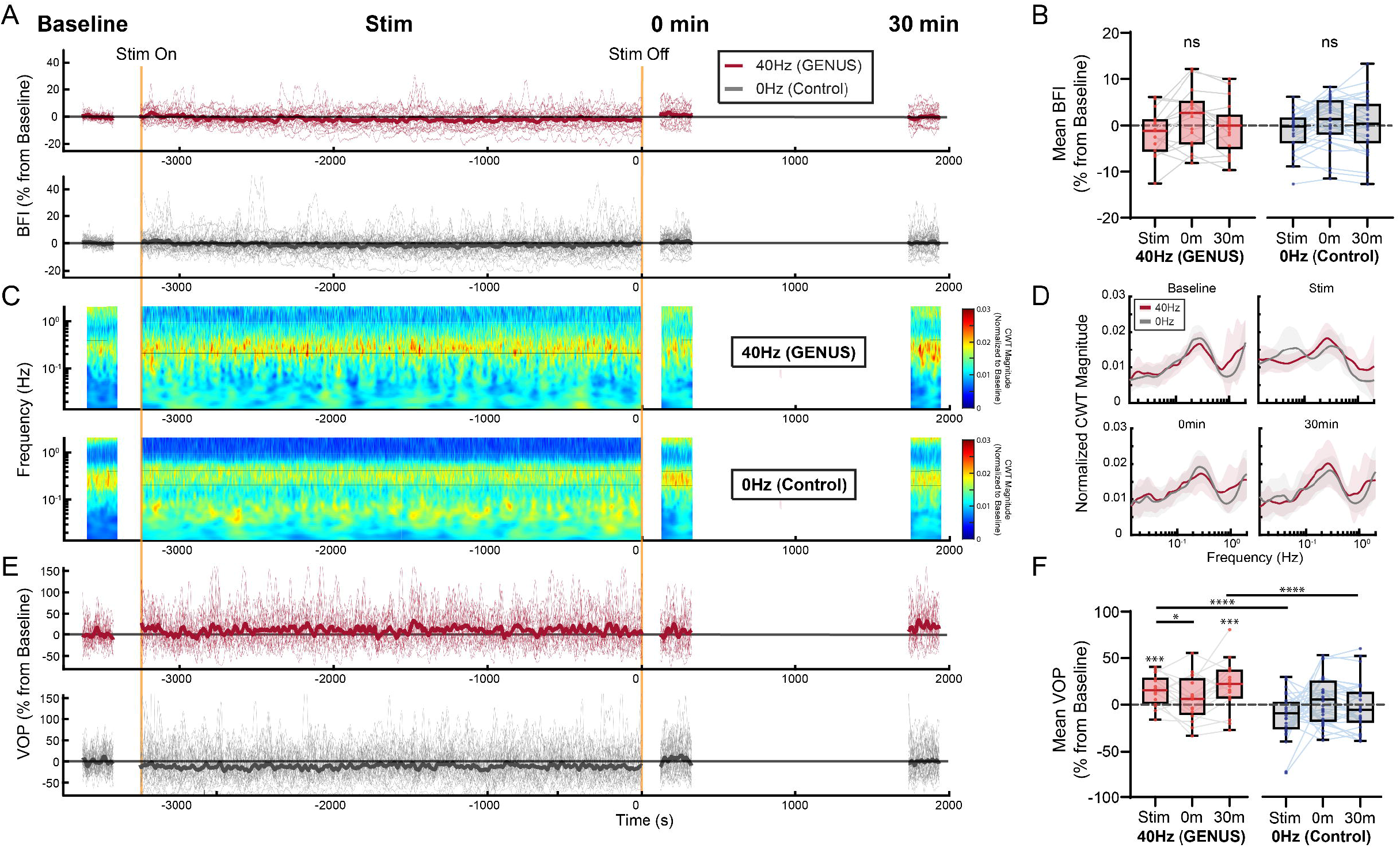
Low-frequency oscillations in blood flow are increased after GENUS stimulation. (A) Spaghetti plot of blood flow index (BFI) for n=18 40Hz GENUS (red) and n=32 0Hz control (gray) datasets, as percent change from Baseline. Group means are plotted in opaque solid lines, while individual time series are plotted in thin transparent lines. (B) Boxplots of percent change in BFI relative to the baseline period for each timepoint. (C) Group average magnitude of the continuous wavelet transform (CWT) of the BFI, normalized to Baseline, for GENUS (n=18) and Control groups (n=32). (D) Spectra calculated from averaging the CWT over each recording time period, with GENUS (red) and Control (gray). Shaded regions show the standard deviation of each group. (E) Spaghetti plot of vascular oscillation power (VOP) averaged over the 0.2-0.4Hz band and normalized to baseline, in a similar style to panel A. (F) Boxplots of percent change in VOP relative to Baseline for each timepoint, treated similarly to panel B. The time series in panels A and E were smoothed with a 20-second moving average to improve visibility. Significance in B and F was determined from Satterthwaite-approximated t-tests using estimated marginal means derived from linear mixed-effects models (details in Methods and Figure 3). *p < 0.05 ***p < 0.001 ****p < 0.0001.

**Figure 3.**
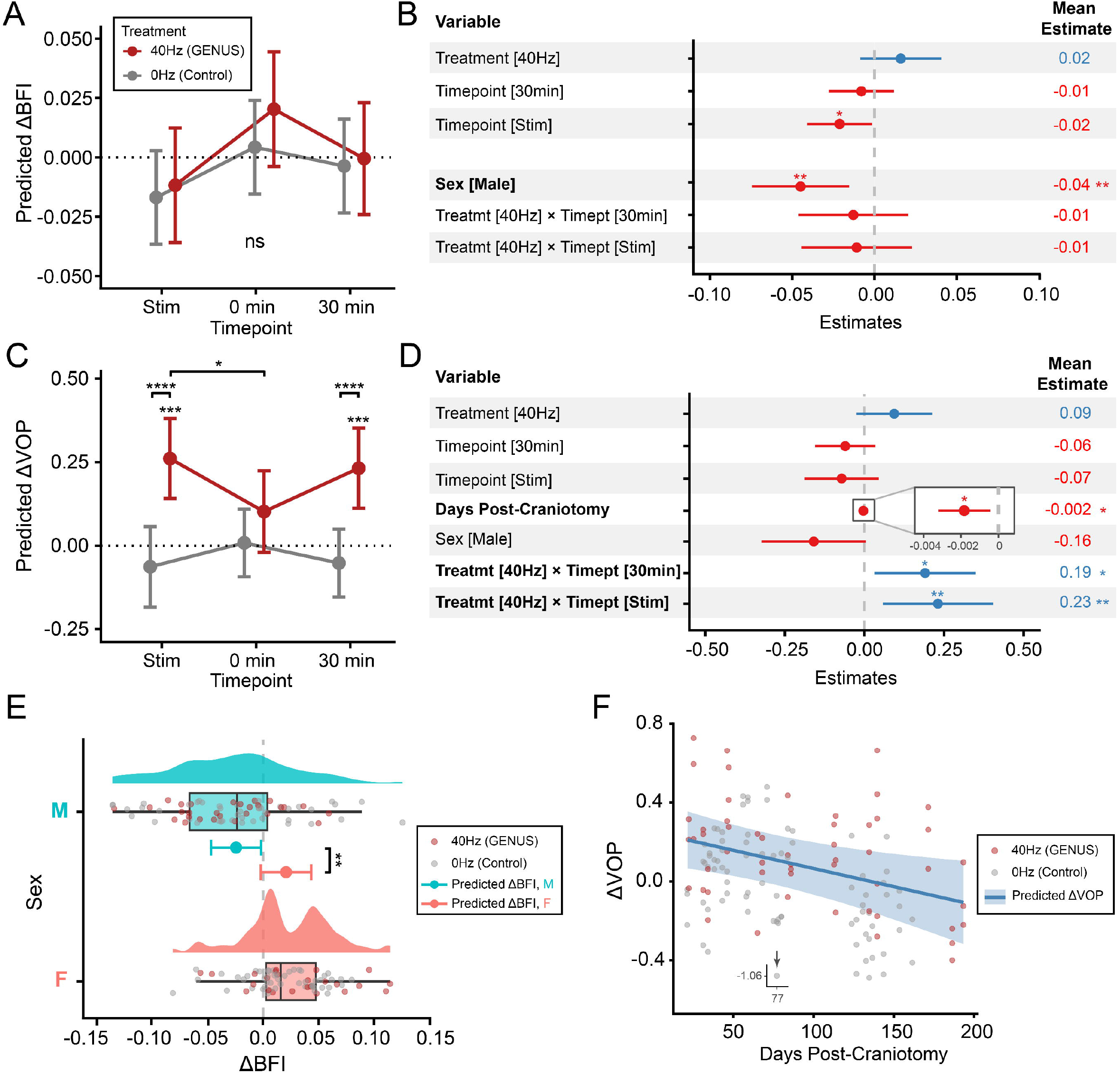
Linear mixed-effects (LME) modeling of GENUS effect on blood flow index (BFI) and vascular oscillation power (VOP). (A) Predicted ΔBFI ∼ (Treatment x Timepoint) + (Sex) + (1|ID) from Model 1. (B) Coefficient estimates from Model 1, with reference levels of Control Treatment, 0min Timepoint and Female Sex. A positive value, marked in blue, indicates a positive effect on ΔBFI, while red is a negative effect. (C) Predicted ΔVOP ∼ (Treatment x Timepoint) + (CraniotomyAge) + (Sex) + (1|ID) from Model 2. (D) Coefficient estimates from Model 2 with similar reference levels to B. (E) Raincloud plot of ΔBFI separated by sex. Points represent ΔBFI data color-coded by treatment. (F) Relationship between days post-craniotomy and predicted ΔVOP. Points represent ΔVOP data color-coded by treatment; the regression line is marked in blue with shaded 95% confidence intervals. An inset shows a single data point that was outside of the range of the y-axis. In all plots, error bars represent 95% confidence intervals. Significance is calculated from t-tests on estimated marginal means with Satterthwaite approximation of degrees of freedom and Tukey adjustment for multiple comparisons. Please see Supplementary Tables ST6, ST7, and ST8 for the summary of tests performed. *p < 0.05, **p < 0.01.

The CWT was computed for each BFI time series, normalized by the mean at baseline for each session, and averaged for GENUS or Control groups (Figure 2C). Strong oscillations in the 0.2-0.4Hz band can be seen in all recordings. However, the magnitude of oscillations appeared to increase during Stim and 30min in the GENUS group, while the Control group showed no change (Figure 2C). The spectra were computed by averaging the CWT magnitude over time and confirm the prominent peak at ∼0.25Hz (Figure 2D). As previous studies typically report LFOs centered around 0.1Hz, we examined whether the frequency discrepancy could be a result of using a CWT rather than a fast Fourier transform. However, the peak frequency in the FFT was consistent with the CWT (Supplementary Figure S5). An additional peak around 0.05Hz was observed during the Control stimulation, but it was not further analyzed (Figure 2D).

The normalized CWT for each session was averaged over the 0.2-0.4Hz band to get a time series of the vascular oscillation power (VOP). The spaghetti plot of VOP relative to Baseline mean shows a positive trend in VOP during Stim and 30min after GENUS, and a negative trend during Stim for the Control group (Figure 2E). Finally, the CWT was averaged over both 0.2-0.4Hz and time to get a scalar quantity of VOP (Figure 2F). VOP was significantly increased during GENUS Stim compared to Baseline (p=0.0001, LME model t-test), but this difference disappeared at 0min as VOP decreased significantly compared to Stim (p=0.04, LME model t-test). To assess changes in VOP during the hour of Stim, 200s periods from the first, middle, and last 20 minutes were analyzed. These showed no differences, indicating that VOP rises soon after stimulation onset and remains stable (Supplementary Figure S6). Interestingly, VOP at 30min after GENUS was again significantly higher than Baseline (p=0.0003, LME model t-test). When comparing treatments, VOP during Stim and 30min was significantly greater in the GENUS group than the Control group (p<0.0001 and p<0.0001 respectively, LME model t-test). We considered whether behavioral alterations could explain the differences between GENUS and Control. However, the percent of time spent whisking during the recording was not different between the groups (Supplementary Figure S7). In summary, GENUS robustly increased VOP during Stim and 30min after, compared to both Baseline and the Control group.

We sought to quantify the effect size of changes in VOP as a result of GENUS. Given the possible influence of many factors including age and sex on BFI and VOP, we used linear mixed-effects models to provide a robust statistical analysis of effects. First, normality of the BFI and VOP data was improved using a log transformation (see Methods), and response variables were calculated as 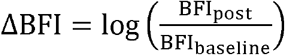 and 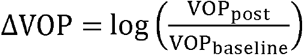. Separate random-intercept models were developed for ΔBFI and ΔVOP (see Methods). Predicted ΔBFI by Model 1 was not significantly different between treatments nor at any timepoint relative to Baseline (LME model t-test) (Figure 3A). However, sex was found to have a significant negative effect of -0.04 on ΔBFI in male mice, which equates to 4% smaller changes in BFI in male mice compared to female mice regardless of treatment or timepoint (Figure 3B,E).

Despite no significant changes in ΔBFI, ΔVOP predicted by Model 2 (see Methods) was significantly increased by 0.26 (95% CI [0.14, 0.38]) during GENUS Stim, equating to a 30% (95% CI [15%, 46%]) increase in VOP compared to Baseline (p=0.0001, LME model t-test) (Figure 3C). This ΔVOP was 0.32 (95% CI [0.14, 0.38]), or 38% (95% CI [22%, 57%]) higher than that of the Control group during Stim (p<0.0001, LME model t-test). However, ΔVOP significantly dropped by -0.16 (95% CI [-0.31, -0.0055]), or 15% (95% CI [0.5%, 27%]), between GENUS Stim and 0min (p=0.04, LME model t-test). At 30min after GENUS, ΔVOP increased again to 0.23 (95% CI [0.11, 0.35]), or 26% (95% CI [12%, 42%]), higher than Baseline (p=0.0003, LME model t-test) and 0.28 (95% CI [0.17, 0.40]), or 33% (95% CI [18%, 49%]), higher than that of the Control group at 30min (p<0.0001, LME model t-test) (Figure 3C). This confirmed the prolonged effect of GENUS on VOP while controlling for multiple variables.

Interestingly, the number of days post-craniotomy significantly decreased the predicted ΔVOP regardless of treatment, by a factor of -0.002, or 0.2%, per day (p=0.01, LME model t-test) (Figure 3D,F). Male mice tended to have 15% lower VOP than female mice, but this was not significant (p=0.07, LME model t-test). A breakdown by sex is available in Supplementary Figure S8. Additionally, the random intercepts showed that individual mice could have response tendencies higher or lower than the group mean, but none were considered outliers (Supplementary Figure S9).

We then sought to identify any spatial patterns in the effect of GENUS or control stimulation across cortical regions within the cranial window. First, BFI maps were calculated by averaging the BFI time series at each pixel. Scalar VOP was computed in a similar way as Figure 2 for each pixel to produce maps of VOP. These maps were registered to the Allen Atlas common coordinates [35] and normalized by the baseline maps before being averaged per treatment group (n=18 datasets for GENUS and n=32 datasets for Control). The BFI maps showed similar patterns when comparing GENUS and Control (Figure 4A). Specifically, blood flow decreased in the posterior part of the window in both groups during stimulation. This effect is slightly alleviated at 0min but returns at 30min. The VOP maps show distinct patterns from BFI (Figure 4B). During stimulation, the medial parts of the windows have increased VOP and at 0min, there is a uniform increase across the window in both groups. However, the Control returns to Baseline levels by 30min, while the VOP of the GENUS group increases further, uniformly across the window (Figure 4B).

**Figure 4.**
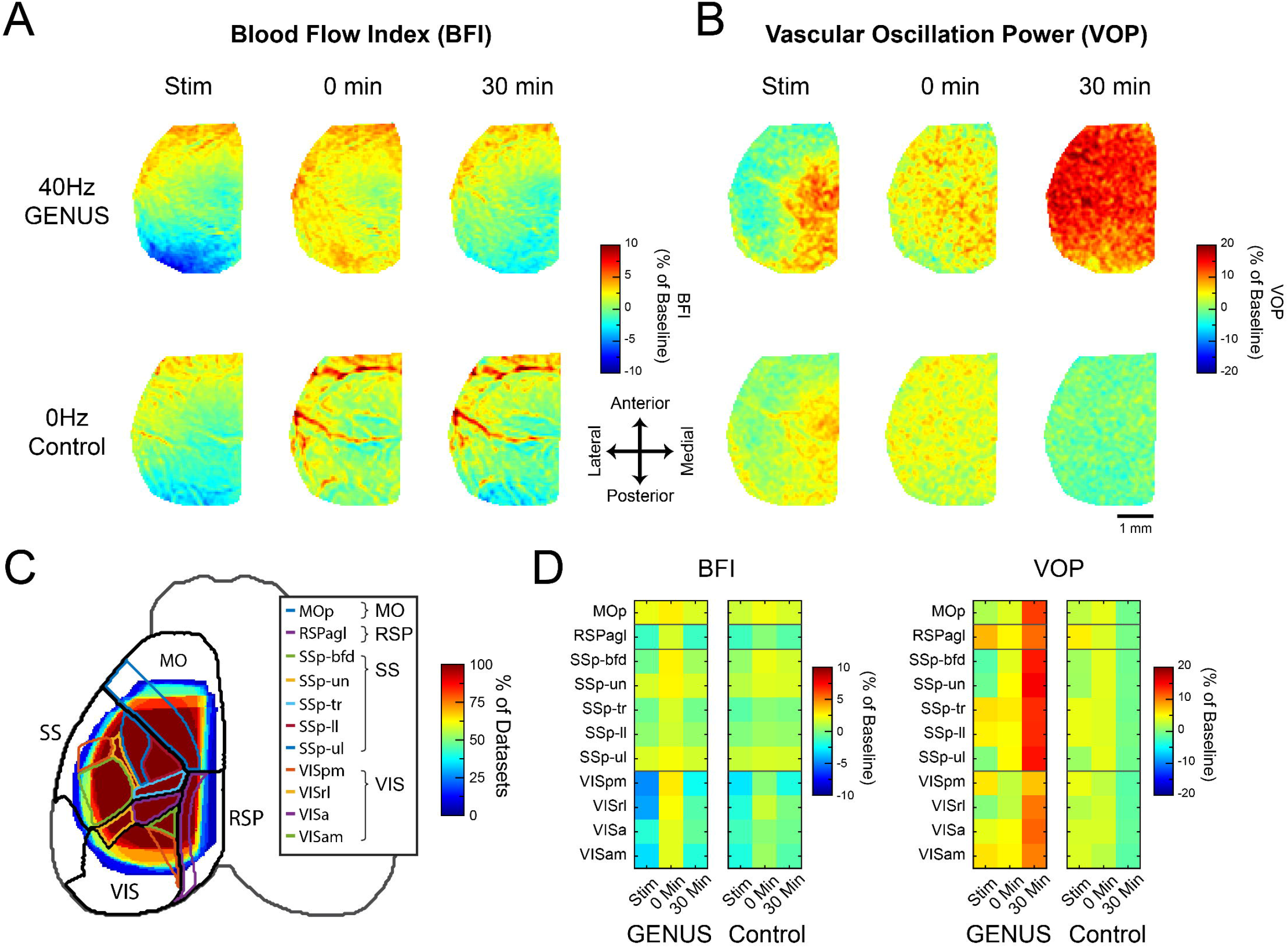
Increases in vascular oscillation power (VOP) span multiple brain regions 30 minutes after GENUS stimulation. Spatial maps of group-average percent change from baseline in (A) blood flow index (BFI) and (B) VOP for 40Hz GENUS (top row) and 0Hz Control (bottom row) during and after stimulation. (C) Heatmap of the overlap of all cranial windows, overlaid with Allen Atlas brain regions. Colored borders are used for regions analyzed in the parcellation. Somatomotor areas (MO); Somatosensory area (SS); Retrosplenial areas (RSP); Visual areas (VIS). A glossary of region names is provided in Supplementary Table ST8. (D) Average BFI and VOP change from baseline for parcels measured under the window. The percentage of datasets for each parcel and timepoint is shown in Supplementary Figure S10. Scale bar: 1mm.

To ascertain the regions where BFI and VOP changes occur, the window was parcellated using the Allen Atlas [35] and the BFI and VOP were averaged over each brain region for each group (Figure 4C). During stimulation, GENUS showed a decrease in blood flow in both the posteromedial area of the visual cortex (VISpm) and the agranular part of the retrosplenial cortex (RSPagl) (Figure 4D). Control stimulation had a slight decrease in the same areas. The decrease in blood flow may indicate that the activated visual region lies outside the cranial window, as the periphery of that region may show a decrease in activity [36]. At 0min, blood flow in all areas was slightly elevated, but the same areas which had decreased flow during Stim also had decreased flow at 30min. Interestingly, the RSPagl and parts of the visual and somatosensory areas had increases in VOP during both GENUS and control stimulation. At 0min, all brain regions were uniformly increased slightly (Figure 3D). Finally, Control showed a return to baseline levels in all regions, while GENUS had elevated VOP in all regions, but slightly less in visual areas. These results show that GENUS stimulation affects VOP in multiple brain regions at 30min, despite the observation that GENUS and Control stimuli engage similar parts of the visual and retrosplenial areas during stimulation.

In a subset of mice, we measured an additional timepoint at 24h after stimulation to assess the return to Baseline levels (GENUS n=11; Control n=23). Figure 5A shows the BFI and VOP at 24h relative to Baseline for both GENUS and Control. We analyzed the data using a separate LME model (Model 3; see Methods). The predicted ΔVOP for the GENUS group was significantly increased compared to Baseline (p=0.04, LME model t-test). This effect was also significantly higher than Control (p=0.02, LME model t-test), suggesting a prolonged effect by GENUS (Figure 5B). However, the mean value was lower than that of the 30min timepoint of the previous model, suggesting a gradual return to baseline. We then sought to evaluate whether natural variability in VOP would obscure later measurements where the VOP would be even closer to Baseline. We measured a further subset of n=6 mice beyond 24h, at 24, 72, 96, and 120h after Control stimulation. The standard deviation of the percent change from Baseline was calculated for each timepoint, which showed a gradual increase over time up to 96h, although it was decreased at 120h. Still, by 48h the deviation was 25%, which is larger than the effect even at 24h, of 21%. Therefore, we expect that a return to baseline beyond 24h could not be easily determined given the natural variability of VOP.

**Figure 5.**
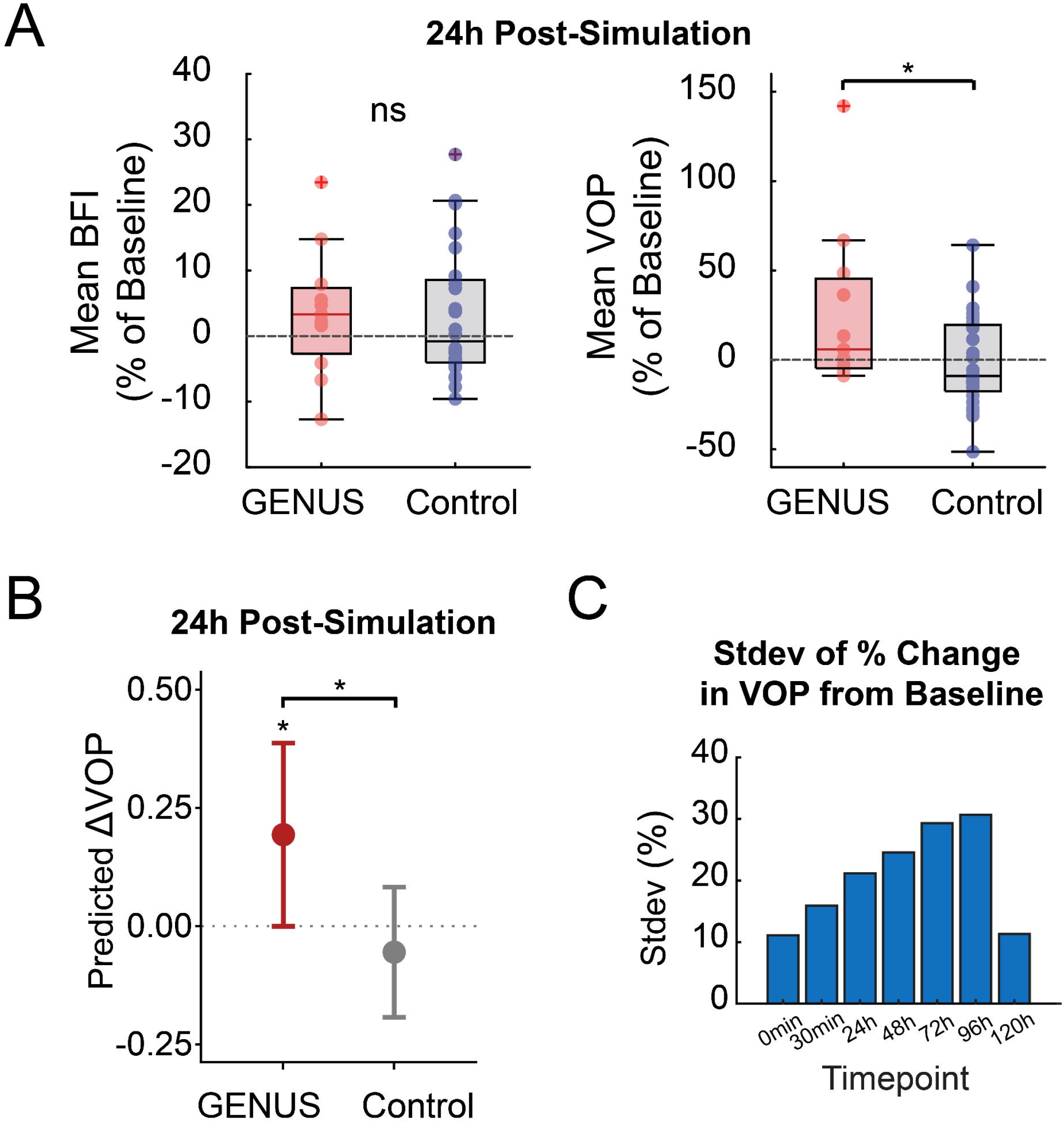
Variability of vascular oscillation power (VOP) over longer timescales obscures return to baseline after GENUS. (A) Boxplots of blood flow index (BFI) (left) and VOP (right). (B) Linear mixed-effects (LME) model comparing predicted ΔVOP ∼ Treatment +(1|ID) from Model 3 between Control (n=23) and GENUS (n=11) groups (p=0.02, LME model t-test). (C) Variability of percent change in VOP in n=6 mice up to 5 days after Control stimulation. Variability rises with time and is expected to obscure the relaxation of VOP to baseline. *p < 0.05

We additionally explored whether alterations of brain state could be associated with changes in VOP. In early pilot experiments, isoflurane anesthesia was used prior to imaging to head-fix the mouse (Supplementary Figure S11A). To assess whether this could have confounded previous results due to altered brain state, we measured BFI continuously before, during and up to 50 minutes after induction by isoflurane (Supplementary Figure S11B). As expected, blood flow increases during induction due to massive dilation of vessels (Supplementary Figure S11C). However, BFI and VOP decrease and remain below baseline up to 30 minutes during recovery. As isoflurane is known to alter neural excitability and brain state, these results suggest that confounding factors related to brain state should be considered while interpreting effects of GENUS on VOP.

## Discussion

In this study, we investigated the effects of 40 Hz visual gamma stimulation (GENUS) on low-frequency oscillations (LFOs; 0.2–0.4 Hz) in blood flow measured using laser speckle contrast imaging (LSCI) in awake mice. Across datasets, vascular oscillation power (VOP) increased during and after GENUS stimulation relative to both pre-stimulation and a 0Hz constant-light control group. Importantly, changes in mean blood flow index (BFI) were modest compared to changes in oscillatory power, indicating that GENUS primarily enhanced the amplitude of intrinsic vascular oscillations rather than producing sustained increases in baseline perfusion. Thirty minutes post-stimulation, the enhancement in VOP was spatially widespread across the cranial window, suggesting global modulation of cortical vascular dynamics rather than a being localized to sensory areas. The treatment effect maintained statistical significance in a linear mixed-effects (LME) model accounting for repeated measures, sex, and days post-craniotomy surgery. Additionally, GENUS-enhanced VOP persisted up to 24 hours post-stimulation. Together, these findings indicate that 40 Hz visual stimulation modulates intrinsic low-frequency vascular dynamics in the cortex, producing effects that outlast the stimulation period.

While prior studies typically show a lower peak frequency in vasomotion closer to ∼0.1Hz [6,21], our data showed a peak frequency of 0.25Hz consistently across mice independent of analysis method. While this lies in the range of vasomotion frequencies shown before [8,18,24,31], more studies are needed to determine the cause of frequency differences and whether they may be influenced by factors such as animal preparation. However, both groups in the present study exhibited similar peaks around ∼0.25Hz, suggesting that the peak shift does not affect the main findings regarding the effect of stimulation.

Notably, our results suggest that intrinsic variability should be considered when interpreting longitudinal measurements. GENUS-treated animals exhibited elevated VOP at 24 hours post-stimulation relative to control. However, measurements were not extended beyond 24 hours as longitudinal analysis revealed that spontaneous vascular oscillations exhibit substantial intrinsic variability across 24–72 hour intervals even in the absence of stimulation. LFOs are known to reflect different myogenic and neuro-modulatory processes that are sensitive to arousal state, autonomic tone, and circadian modulation [37]. In awake animals, even subtle changes in behavioral state or baseline cortical excitability can significantly influence spontaneous hemodynamic oscillations [38]. Thus, while our findings demonstrate persistence of GENUS-associated modulation at 24 hours, inherent variability limits the measurement of small effect sizes at longitudinal timepoints.

The dependence of LFOs on factors such as behavioral state may also explain absence of a clear effect at the immediate 0min post-stimulation timepoint. The transition from light to dark conditions at the end of the stimulus period could influence the mouse’s arousal or brain state, masking differences between treatments. While whisker recordings did not indicate a change in behavior at 0min compared to baseline in both groups, a more sensitive measure for arousal and brain state, such as pupil diameter, could elucidate differences. By 30min post-stimulation, the effect of the light-dark transition would have passed and allowed the GENUS-associated increase in VOP to become more apparent. Together, these observations underscore the sensitivity of spontaneous vascular oscillations to subtle behavioral state changes and highlight the importance of accounting for state when interpreting longitudinal vascular measurements.

Additionally, spectral analysis revealed a prominent increase in very-low-frequency power (∼0.05 Hz) during the constant-light condition. Prior work demonstrates that very-low-frequency vasomotor oscillations dominate during low-arousal states, including wakefulness and non-REM sleep [39,40]. In contrast, 40 Hz stimulation was associated with no increase in this very-low-frequency oscillatory regime and an increase in the low-frequency (∼0.25 Hz) fluctuations, consistent with a shift toward a more aroused autonomic state [39]. Together, these observations could suggest that constant light permits the intrinsic very-low-frequency vasomotion characteristic of a quiescent regulatory state, whereas 40 Hz stimulation shifts toward an arousal-dominated regime that suppresses slow vasomotor coherence.

These observations may link GENUS to specific pathways involving neuromodulators such as norepinephrine, which controls arousal and drives widespread LFOs in sleep [7,8]. GENUS has already been linked to sleep and neuromodulation via increased extracellular adenosine post-stimulation, which promotes sleep via a buildup of sleep pressure and is tied to increased LFOs and glymphatic flow [29,41]. This extracellular adenosine increase is likely caused by the entrained gamma activity, where demanding high-frequency neural activity causes adenosine nucleotide release that is catabolized into adenosine [42]. This adenosine could inhibit vasoactive arousal-related modulators, such as basal forebrain acetylcholine and locus coeruleus norepinephrine [42,43], and modify their concentrations or dynamics. However, it remains to be determined whether these LFO dynamics could be driven by acetylcholine or norepinephrine dynamics or whether the oscillations could simply be intrinsically produced after elevation or depletion of a neuromodulator input.

Spatial analysis revealed that GENUS-induced increases in VOP were broadly distributed across the imaged cortex, yet regionally heterogeneous in magnitude. VOP was initially increased in visual and retrosplenial areas during stimulation before expanding to the entire imaged cortex by 30min afterward. While modulation of visual cortex was expected due to the use of visual-only stimulation, it is possible that multisensory stimulation could increase VOP during stimulation in those associated regions as well. The distributed effect at 30min could suggest propagation of prolonged gamma-driven activity through distributed cortical networks rather than a purely local sensory-evoked response.

Several limitations of the present study should be acknowledged. First, only 40 Hz stimulation was tested, and frequency specificity was not systematically evaluated. Second, time of day was not controlled across experimental sessions, which may have contributed to variability in oscillatory magnitude due to circadian and state-dependent factors on vascular dynamics. Incorporating multiple indicators of arousal besides whisking would allow more precise dissection of state-dependent contributions to vascular oscillations. Importantly, electrophysiological recordings were not obtained in the present study. As a result, although 40 Hz visual stimulation is widely reported to entrain cortical gamma activity [25,41], we cannot directly determine whether the prolonged LFOs are coupled by prolonged effects on gamma-band activity. Future studies combining simultaneous neural and vascular measurements will be necessary to test whether GENUS improves vascular oscillatory dynamics through direct modulation of gamma activity and neurovascular coupling mechanisms.

Age-related factors also limited the study by increasing variability of VOP. The LME modeling uncovered a significant negative relationship between VOP magnitude and days post-craniotomy. Chronic cranial window preparations are known to induce gradual alterations in cortical physiology, including low-grade inflammation, glial activation, and remodeling of vasculature such as the growth of collateral arteries [32,44]. Such factors could contribute to the observed decline in VOP over time. However, while this effect may increase variability in VOP, it was observed equally in both GENUS and control conditions and does not affect the main findings of the study.

In summary, the present findings demonstrate that 40 Hz visual stimulation produces prolonged and widespread enhancement of low-frequency vascular oscillations in the cerebral cortex of mice. Future studies could further link GENUS-modulated vascular dynamics to glymphatic flow and clearance to support previous research using GENUS to enhance clearance in Alzheimer’s disease [25,26]. Such a link would provide mechanistic rationale behind exploring GENUS as a therapeutic in other diseases of vascular or glymphatic dysfunction, such as ischemic stroke. Optogenetic gamma stimulation of GABAergic neurons post-stroke has already shown improvements in tissue and behavioral outcomes [45]. In parallel, these results could be applied to human studies, as LFOs can be measured using noninvasive modalities such as functional near-infrared spectroscopy (fNIRS) or resting-state functional magnetic resonance imaging (fMRI). Determining whether modulation of vascular oscillations contributes to reported cognitive or pathological improvements will be critical for establishing whether GENUS holds therapeutic promise for disorders involving neurovascular dysfunction.

## Acknowledgements

This work was supported by the National Institutes of Health (R01NS127156 and R01NS122742). We thank the Boston University Master’s in Statistical Practice Consulting program, namely Himani Yadav and Masanao Yajima, for facilitating the development of the statistical analyses. The experimental portion of this study was facilitated by the Boston University Neurophotonics Center.

## Author Contribution Statement

DAB and AD conceived the study. JJ, PRB, and RPT performed animal preparation and data collection. RPT and PRB analyzed the data. BCR and NC developed the stimulation device. RV, ZG, and YS developed the statistical analyses. RPT, PRB and EL created the figures. RPT, PRB, and EL wrote the paper. DAB, KK, ŞEE, and AD provided critical discussion and revision of the article. All authors approved the final version.

## Supplementary Information

Supplemental material for this article is available online.

## Data availability

Data will be made available via DANDI Archive ID 001790.

