## SupplementaryTablesAndFigures for "Visual gamma stimulation causes prolonged enhancement of low-frequency blood flow oscillations across cortical regions in mice"

**Supplementary Figures**

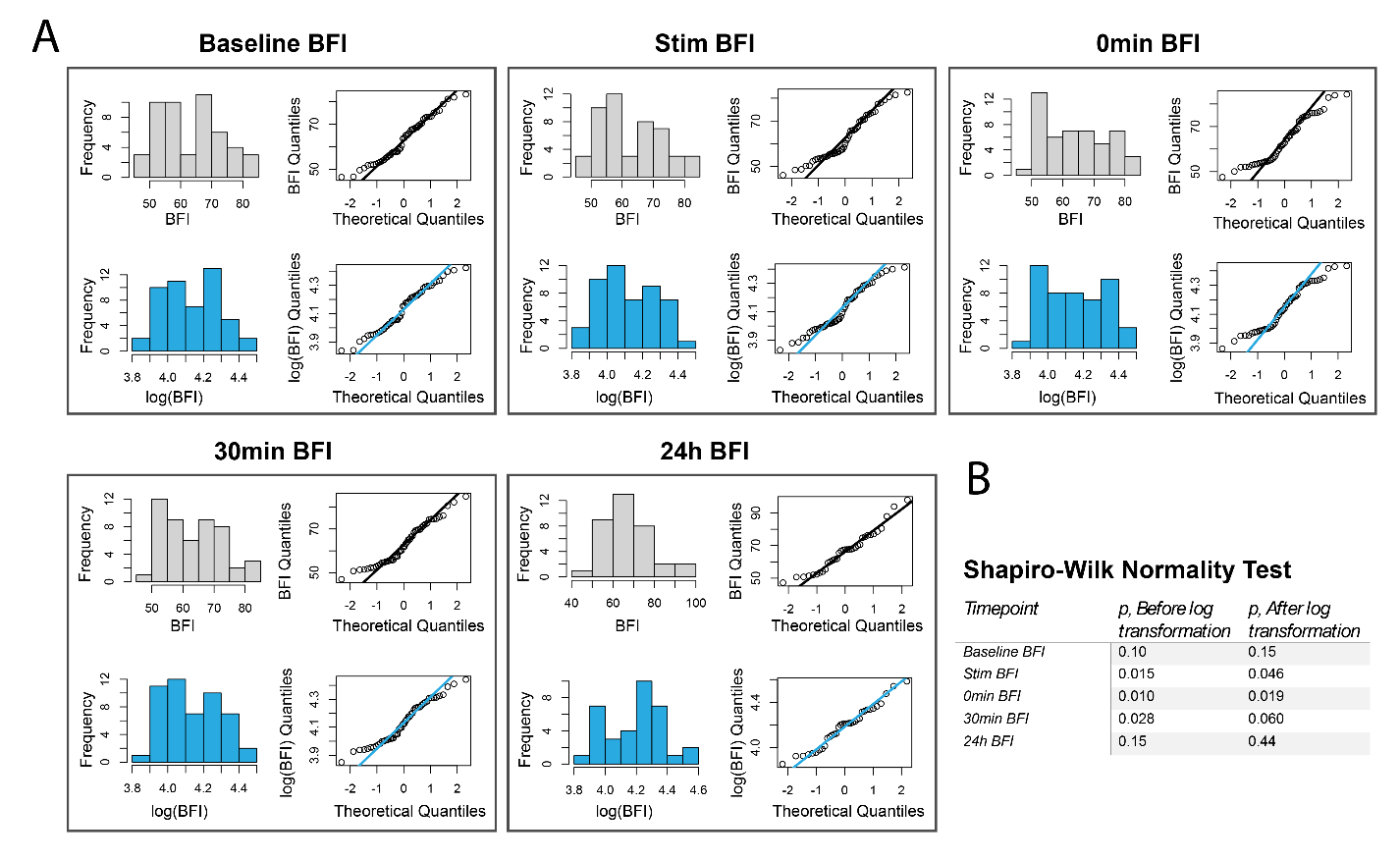

Supplementary Figure S1: A) Log transform of blood flow index (BFI) data. Histograms and QQ plots of BFI (gray) and log(BFI) (blue) show that the log transform improves the normality of the data. B) A Shapiro-Wilk normality test was performed for each timepoint before and after log transformation. While 0min BFI still had p < 0.05, the normality was greatly improved and deemed acceptable for further analysis.

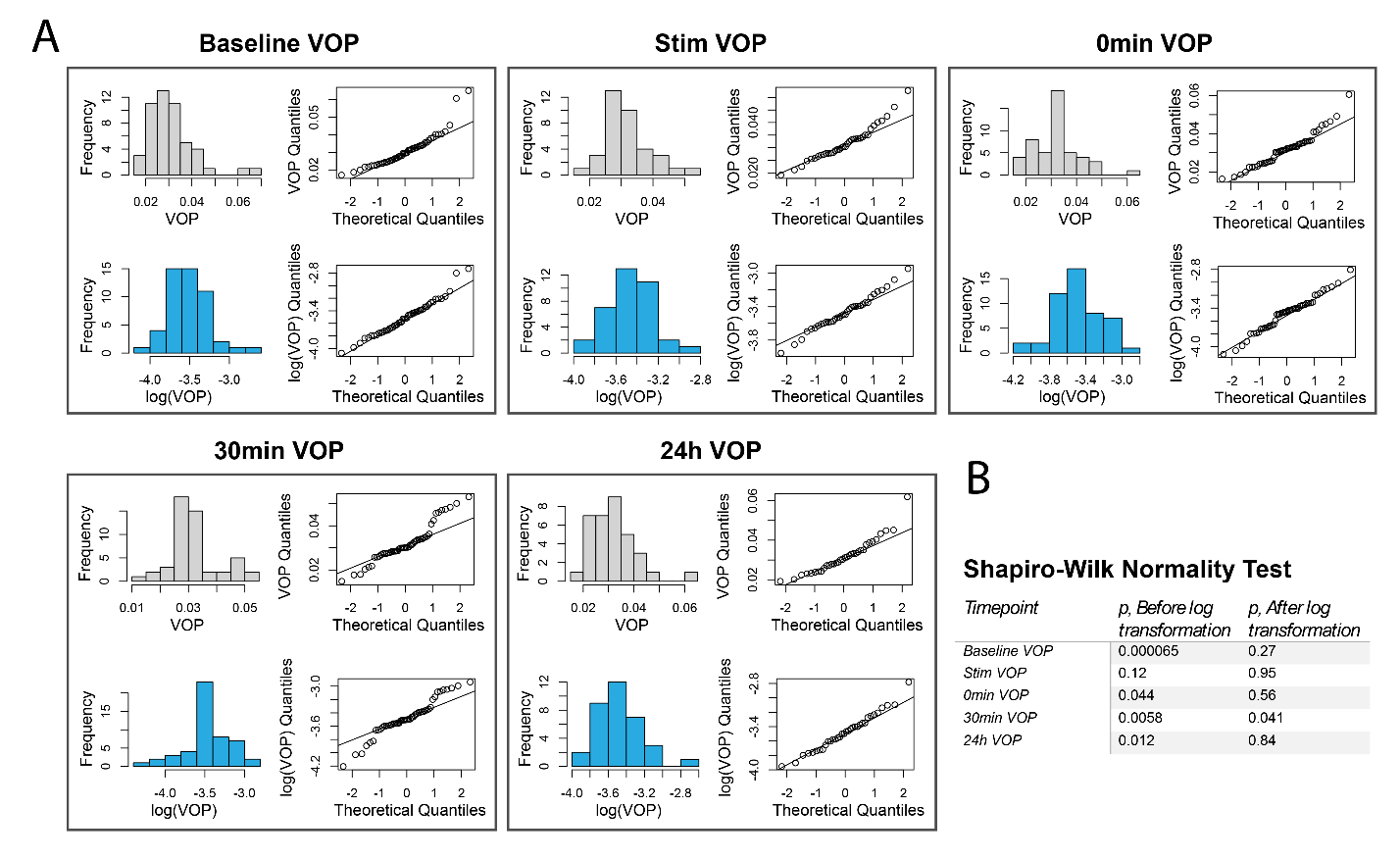

Supplementary Figure S1: A) Log transform of vascular oscillation power (VOP) data. Histograms and QQ plots of VOP (gray) and log(VOP) (blue) show that the log transform improves the normality of the data. B) A Shapiro-Wilk normality test was performed for each timepoint before and after log transformation. While 30min VOP still had p < 0.05, the normality was greatly improved and deemed acceptable for further analysis.

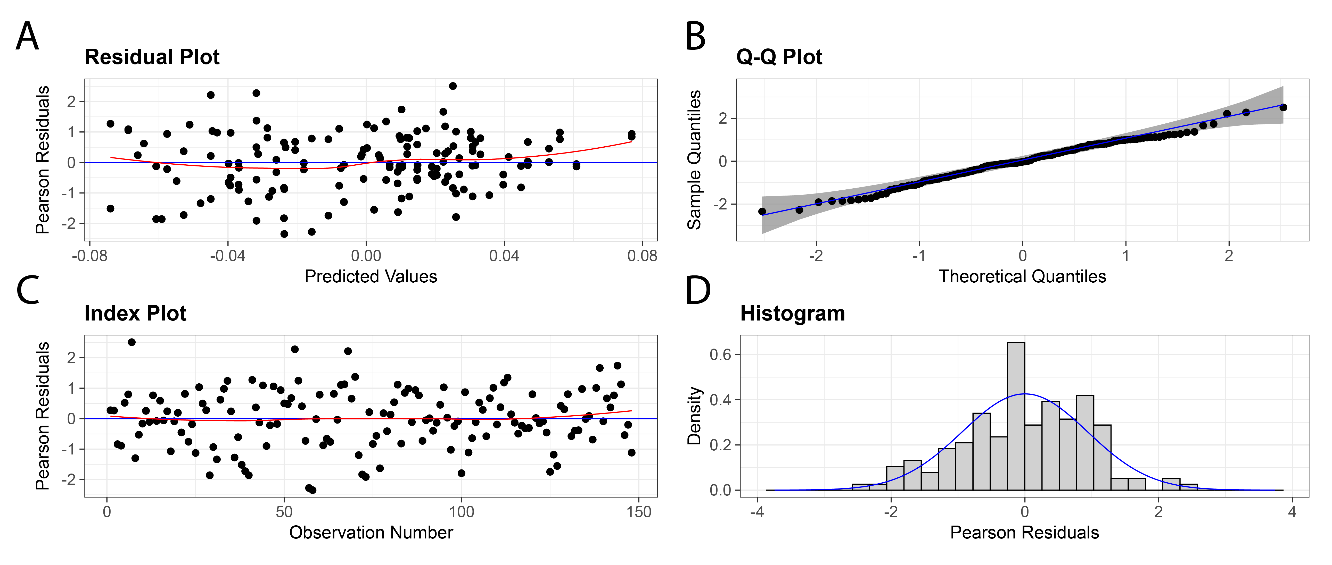

Supplementary Figure S3: Normality, homoscedasticity of linear mixed-effects Model 1. A) Plot of residuals vs. predicted values of $\Delta\mathrm{BFI}$ shows homoscedasticity. B) Q-Q plot shows the normality of residuals. C) Plot of residuals vs. observation number shows few outliers. D) The histogram of residuals further reinforces the normality of residuals.

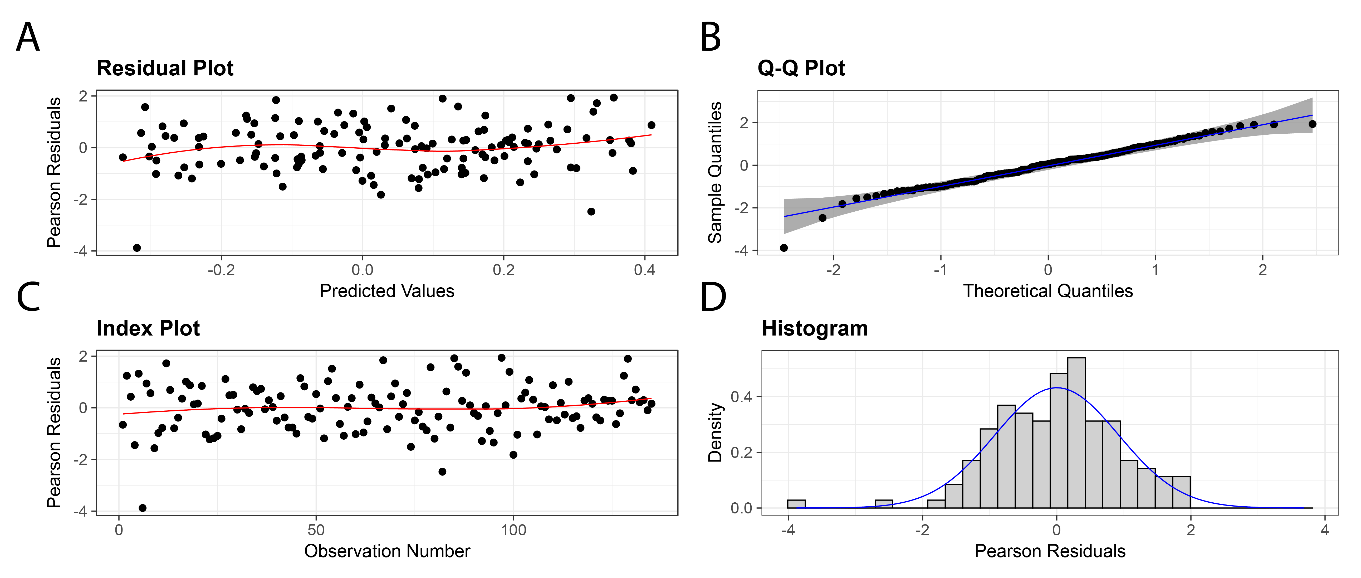

Supplementary Figure S4: Normality, homoscedasticity of linear mixed-effects Model 2. A) Plot of residuals vs. predicted values of $\Delta\mathrm{VOP}$ shows homoscedasticity. B) Q-Q plot shows the normality of residuals. C) Plot of residuals vs. observation number shows few outliers. D) The histogram of residuals further reinforces the normality of residuals.

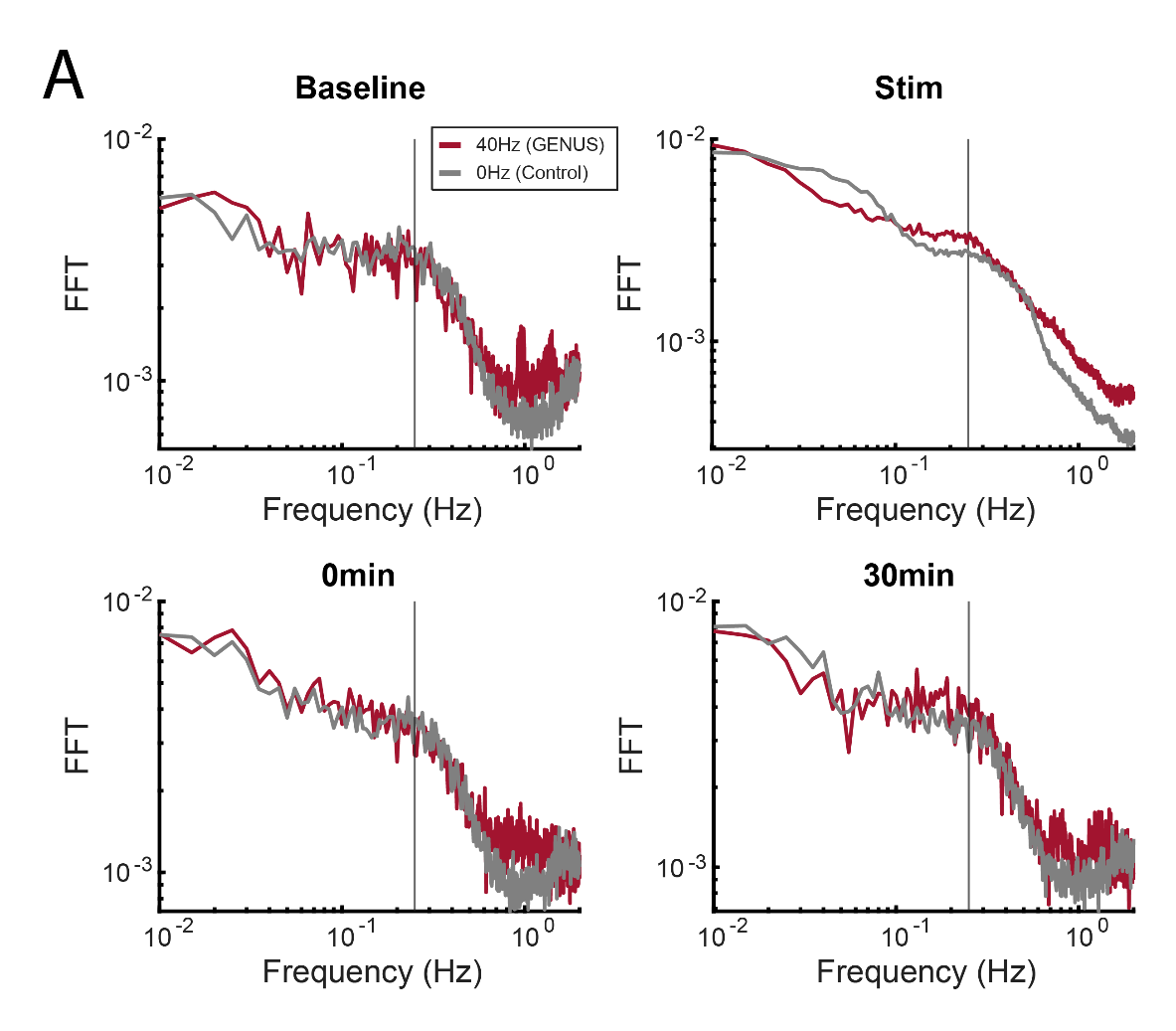

Supplementary Figure S5: A) Average fast Fourier transform (FFT) spectra for each treatment group and timepoint. The peak frequency is around 0.25Hz (vertical line), which is consistent with continuous wavelet transform-computed spectra. The Stim period was chunked into 15 non-overlapping windows of 200s and the FFTs for each period were averaged to get the spectrum.

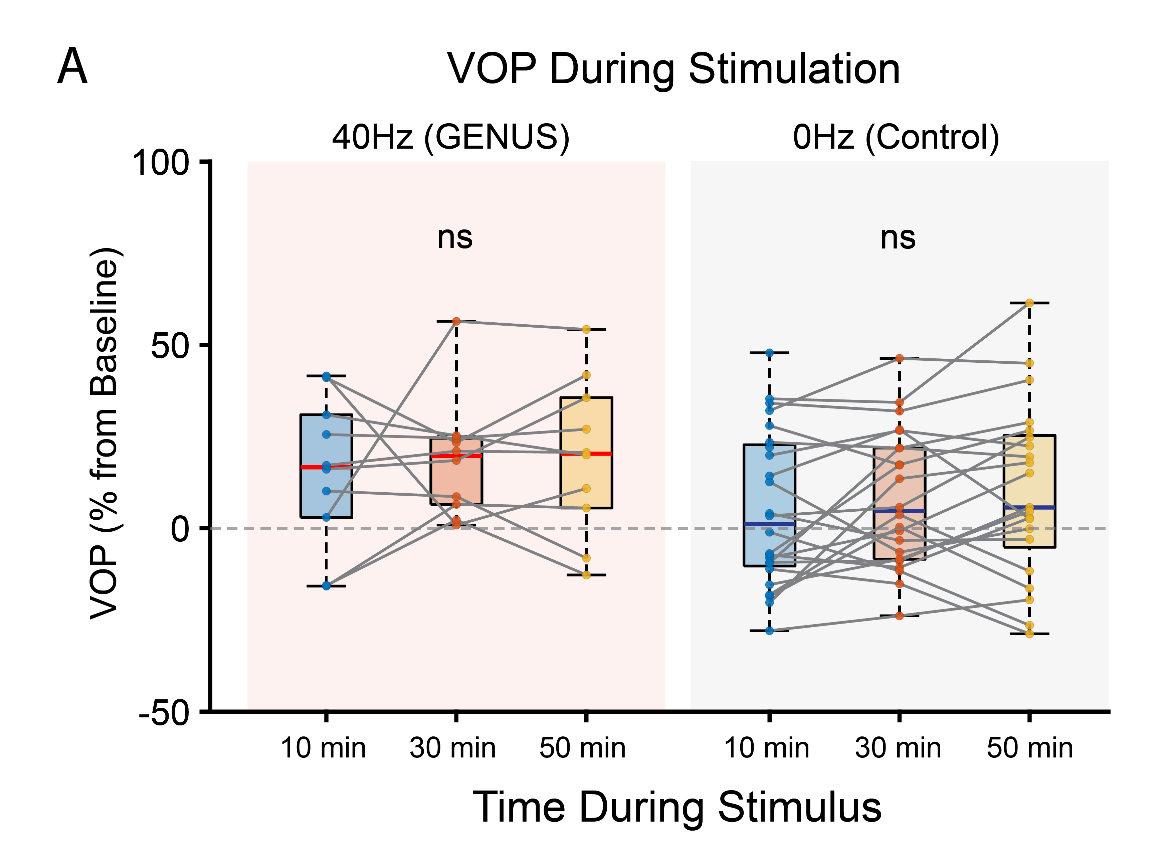

Supplementary Figure S6: A) Vascular oscillation power (VOP) computed for 200s windows during the first, middle and last 20 minutes of the 1hr of stimulation. There were no differences within each treatment (Wilcoxon signed rank test), indicating a quick onset of VOP increase after beginning stimulation.

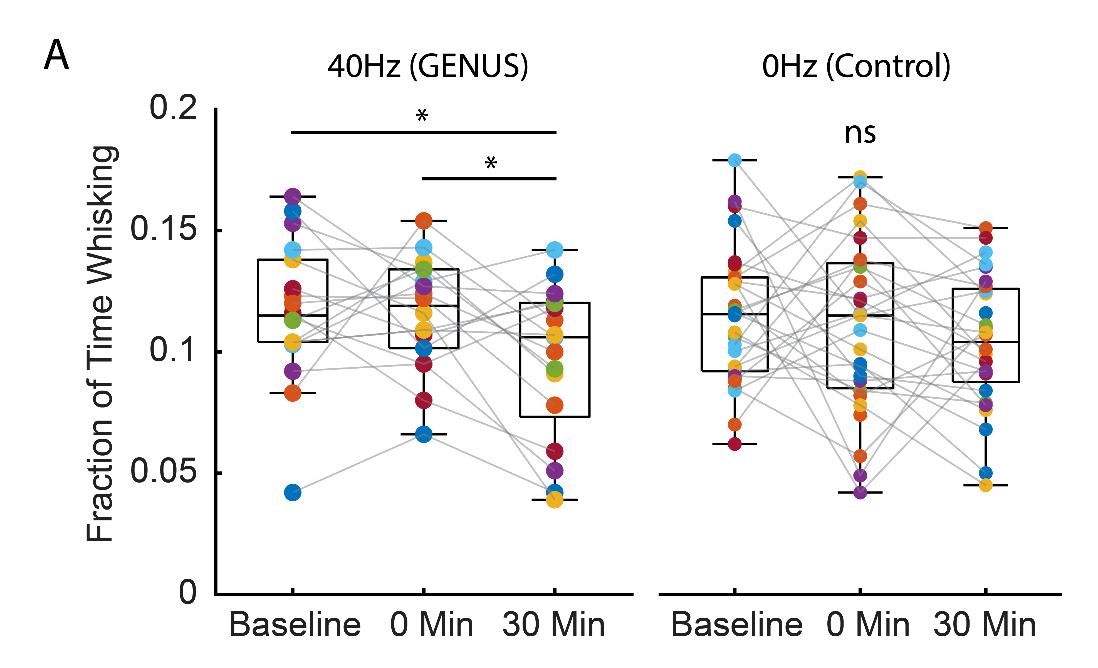

Supplementary Figure S7: A) Fraction of the 200s laser speckle contrast imaging window during which the mouse was whisking. While the GENUS group did show a decrease in whisking at 30min (Wilcoxon signed rank test), there were no differences between the groups (Wilcoxon rank sum test).

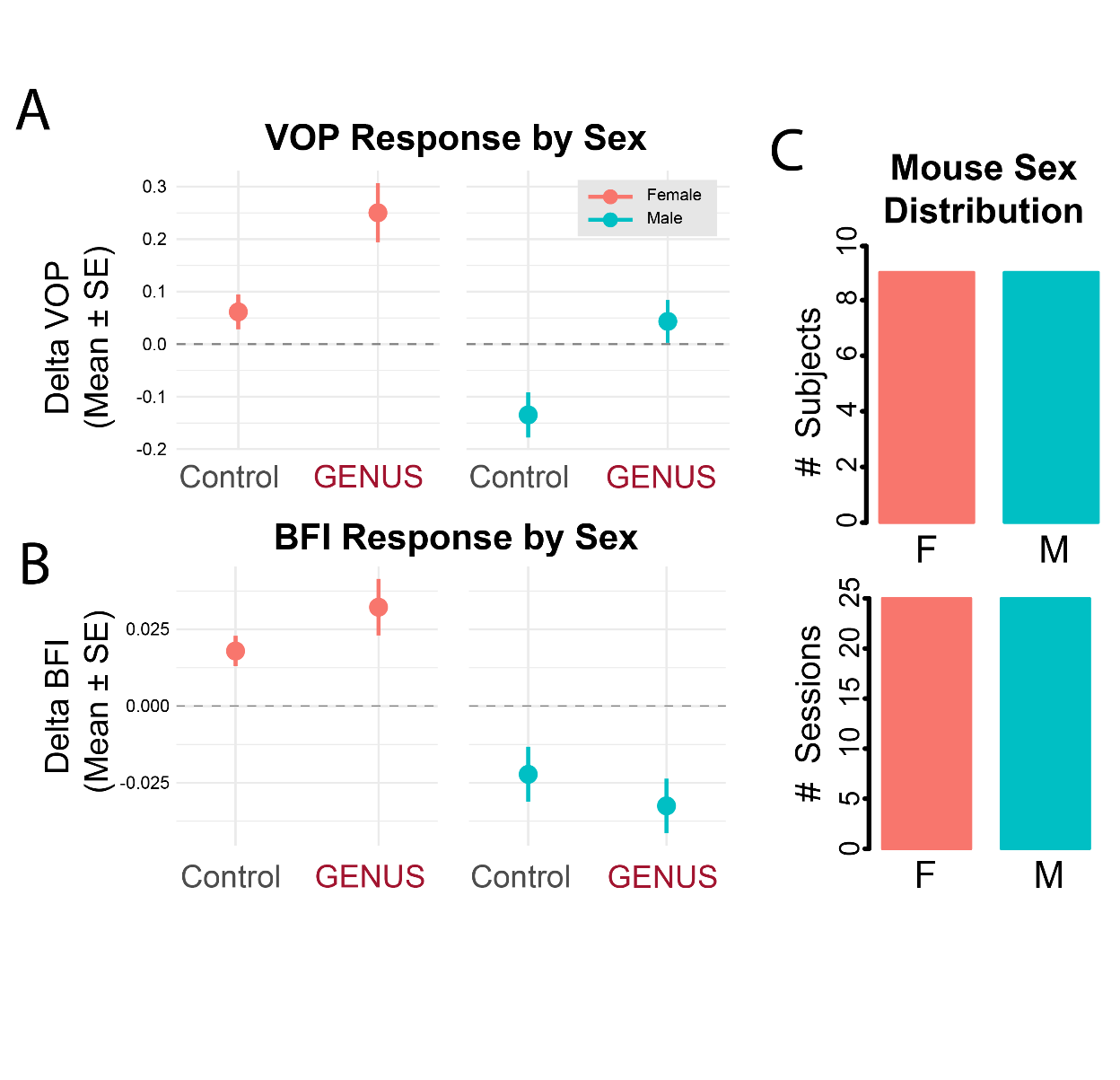

Supplementary Figure S8: Breakdown of datasets by sex. A) While male mice tended to have a decreased $\Delta\mathrm{VOP}$, the GENUS group had higher $\Delta\mathrm{VOP}$ in both sexes. Error bars represent 95% confidence intervals. B) Female mice had a tendency for greater increases in BFI in GENUS compared to Control, while male mice had greater decreases in BFI in the GENUS group. (C) The distribution of sexes was equal in terms of subject numbers (left) and dataset numbers (right).

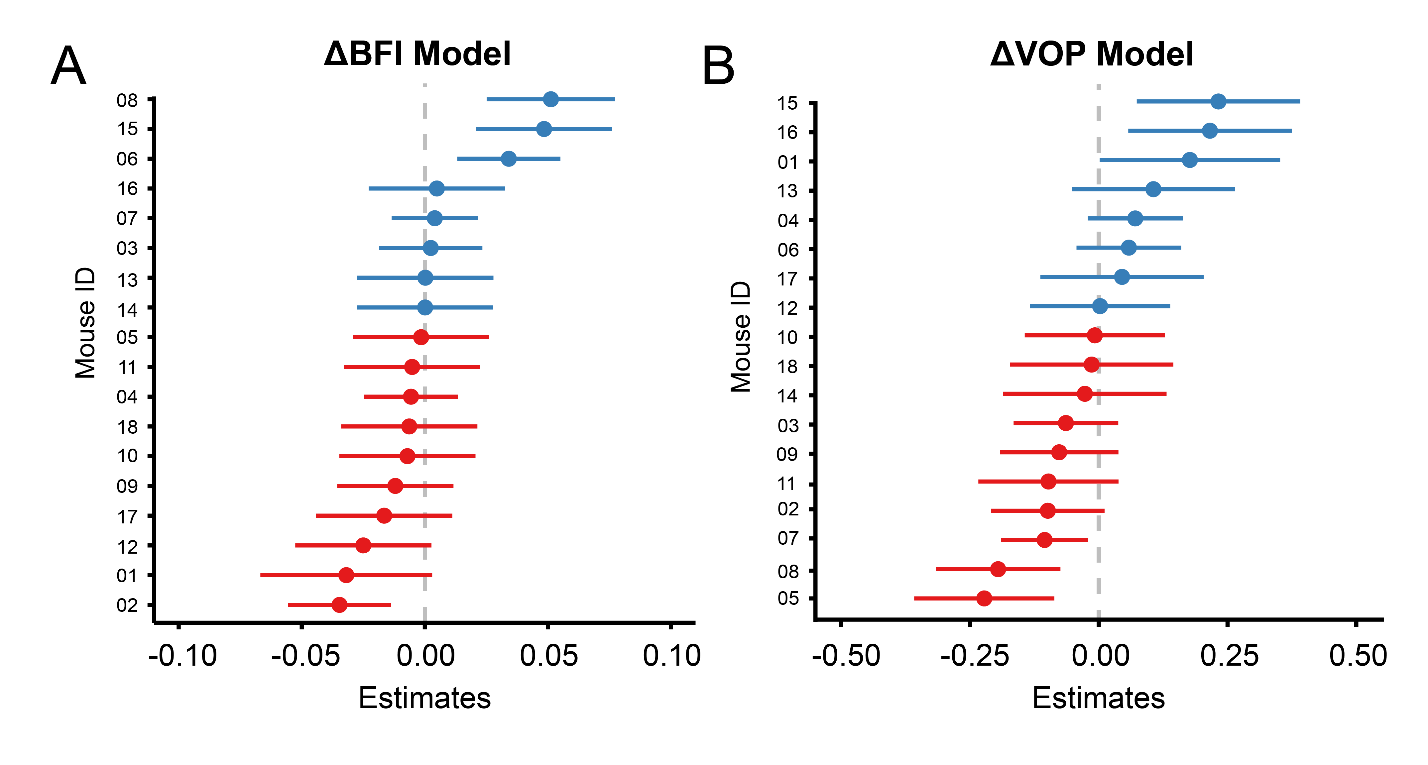

Supplementary Figure S9: Summary of random intercepts of (A) Model 1 and (B) Model 2. Error bars represent 95% confidence interval. Blue color indicates the mouse had a tendency in effect above the group mean, while red indicates a tendency below the group mean.

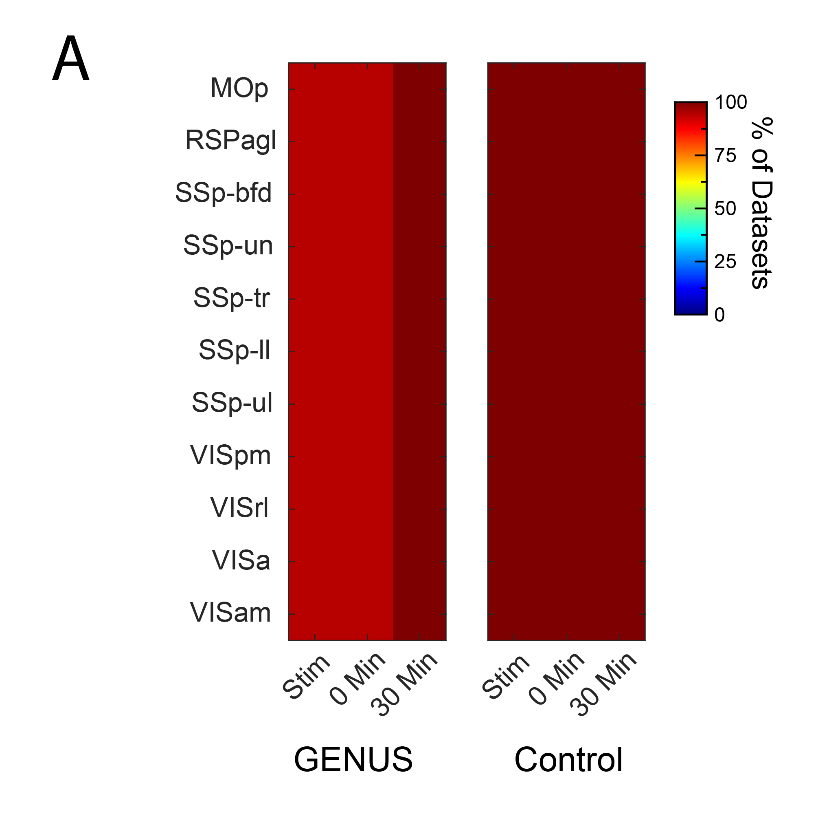

Supplementary Figure S10: A) Percentage of datasets represented in each parcel and each timepoint of Figure 4.

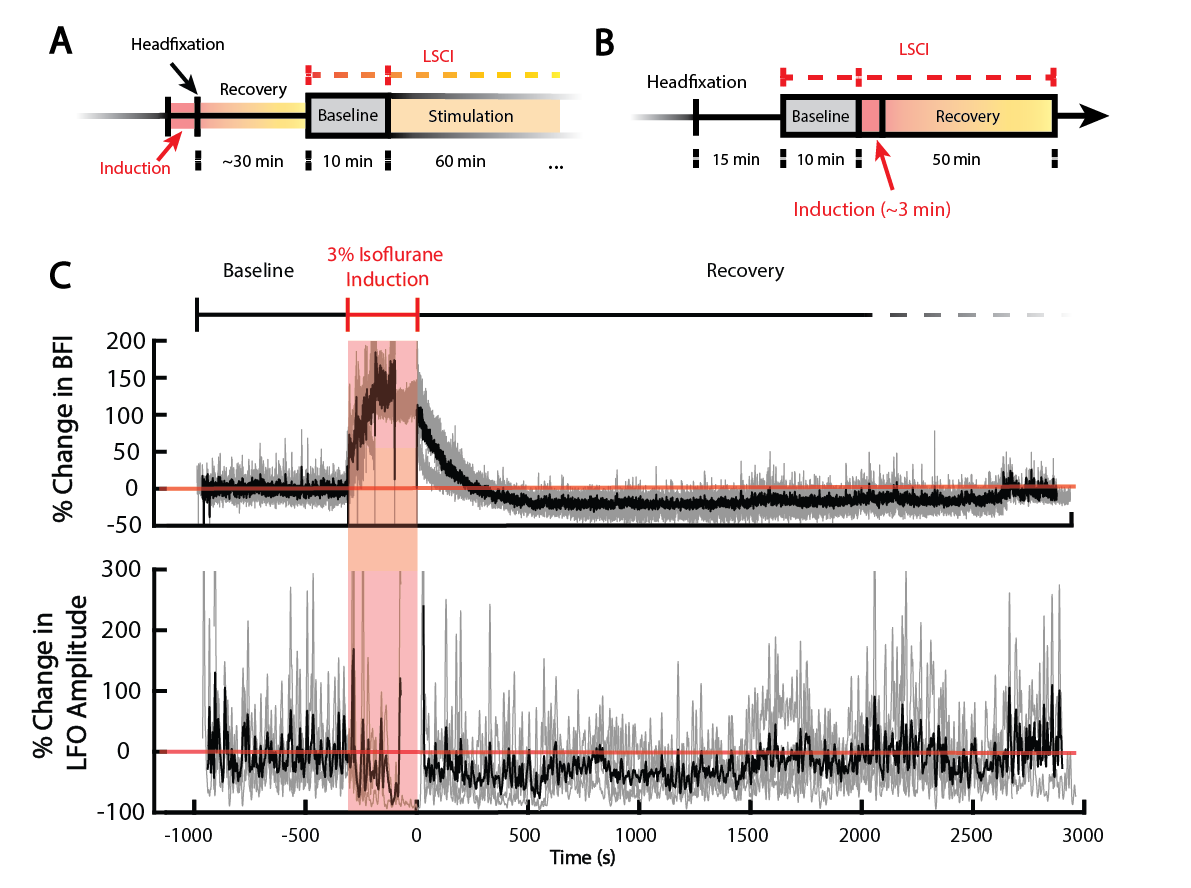

Supplementary Figure S11: Induction by isoflurane anesthesia causes prolonged decreases in blood flow and 0.2-0.4Hz vascular oscillation power (VOP). (A) Timeline for initial pilot experiments which were confounded by isoflurane induction prior to imaging. (B) Timeline for isoflurane induction and recovery experiments. To assess the prolonged effects of isoflurane anesthesia on neurovascular dynamics, 3 mice underwent 1-hour laser speckle contrast imaging sessions using LSCI in the absence of any stimulation. Each session began with a 10-minute baseline recording, followed by administration of 3% isoflurane in 0.8 L/min oxygen via a nose cone. Once a respiratory rate of 80–100 breaths per minute was reached, the isoflurane was discontinued, and the mouse was allowed to recover for the remaining 45–50 minutes while LSCI data continued to be acquired. (C) Spaghetti plots of percent change in BFI (top) and VOP (bottom) relative to baseline period in n=3 mice. Isoflurane causes rapid increases in BFI during induction and a prolonged BFI and VOP deficit during recovery, for up to ~30 minutes.

Supplementary Table ST1: Sample size calculation using anticipated values for % change in vascular oscillation power. Sample size selected is at 80% statistical power and alpha level 0.05.

|  | **Anticipated Values** | | |  | |  | |
| --- | --- | --- | --- | --- | --- | --- | --- |
|  | **Mean (%)** | **Std. Dev (%)** | |  | |  | |
| **Control** | 100 | 20 | |  | |  | |
| **GENUS** | 120 | 20 | |  | |  | |
|  | **Sample Size Needed in Each Group** | | | | | | |
|  |  | **Power** | | | | | |
|  |  | **95%** | **90%** | | **80%** | | **50%** |
| **alpha level** | **0.10** | 22 | 17 | | 12 | | 5 |
|  | **0.05** | 26 | 21 | | **16** | | 8 |
|  | **0.02** | 32 | 26 | | 20 | | 11 |
|  | **0.01** | 36 | 30 | | 23 | | 13 |

Supplementary Table ST2: Pearson correlation between subject factors to analyze collinearity.

| **Factor** | **Mouse Session #** | **Mouse Age (days)** | **Days Post-Craniotomy** |
| --- | --- | --- | --- |
| **Mouse Session #** | 1.00 | 0.67 | 0.61 |
| **Mouse Age (days)** |  | 1.00 | 0.88 |
| **Days Post-Craniotomy** |  |  | 1.00 |

Supplementary Table ST3: Correlated time-related variables (Session #, Age, and Days After Craniotomy) were reduced by selecting the variable which had the most significance and largest influence on model performance. Three models were compared using the formula $\Delta BFI \sim Treatment\times Timepoint+Variable+Sex+(1|ID)$, where $Variable$ was either Session #, Age, or Days After Craniotomy.

| **Variable** | **AIC** | **BIC** | **β** | **p** |
| --- | --- | --- | --- | --- |
| Session # | -692.5 | -653.0 | -0.0010 | 0.73586 |
| Age | -693.2 | -653.8 | -0.000073 | 0.35809 |
| Days After Craniotomy | -692.5 | -653.1 | -0.000044 | 0.66051 |

Supplementary Table ST4: Summary for Model 1: $\Delta BFI \sim Treatment \times Timepoint+Sex+(1|ID)$, with restricted maximum likelihood (REML) estimation and Satterthwaite degrees of freedom. The reference levels are: Control treatment, 0min timepoint, female sex. Data includes 135 observations and 18 subjects (ID). Because the t-tests reported in this table are not adjusted for multiple comparisons, please refer to Supplementary Table ST7 for a comprehensive summary of the p-values with Tukey adjustment.

|  |  | **Std. Dev** | **Variance** | **ICC** |  |  |
| --- | --- | --- | --- | --- | --- | --- |
| **Random Effects** | ID (Intercept) | 0.0279 | 0.000778 | 0.326 |  |  |
|  | Residual | 0.0401 | 0.00161 |  |  |  |
|  |  | **β** | **SE** | **df** | **t** | **p** |
| **Fixed Effects** | (Intercept) | 0.027 | 0.012 | 26.2 | 2.195 | **0.037** |
|  | Treatment [GENUS] | 0.016 | 0.012 | 130. | 1.282 | 0.20 |
|  | Timepoint [30min] | -0.0080 | 0.010 | 124. | -0.794 | 0.42 |
|  | Timepoint [Stim] | -0.021 | 0.010 | 124. | -2.115 | **0.036** |
|  | Sex [M] | -0.045 | 0.015 | 14.9 | -3.004 | **0.0090** |
|  | Treatment [GENUS] x Timepoint [30min] | -0.013 | 0.017 | 124. | -0.760 | 0.45 |
|  | Treatment [GENUS] x Timepoint [Stim] | -0.011 | 0.017 | 124. | -0.637 | 0.52 |

Supplementary Table ST5: Correlated time-related variables (Session #, Age, and Days After Craniotomy) were reduced by selecting the variable which had the most significance and largest influence on model performance. Three models were compared using the formula $\Delta VOP \sim Treatment\times Timepoint+Variable+Sex+(1|ID)$, where $Variable$ was either Session #, Age, or Days After Craniotomy.

| **Variable** | **AIC** | **BIC** | **β** | **p** |
| --- | --- | --- | --- | --- |
| Session # | -18.6 | 10.4 | -0.036 | 0.065 |
| Age | -22.4 | 6.7 | -0.0015 | 0.011 |
| Days After Craniotomy | -22.5 | 6.6 | -0.0018 | **0.0090** |

Supplementary Table ST6: Summary for Model 2: $\Delta VOP \sim Treatment \times Timepoint+CraniotomyAge+Sex+(1|ID)$, with restricted maximum likelihood (REML) estimation and Satterthwaite degrees of freedom. The reference levels are: Control treatment, 0min timepoint, female sex. Data includes 135 observations and 18 subjects (ID). Because the t-tests reported in this table are not adjusted for multiple comparisons, please refer to Supplementary Table ST8 for a comprehensive summary of the p-values with Tukey adjustment.

|  |  | **Std. Dev** | **Variance** | **ICC** |  |  |
| --- | --- | --- | --- | --- | --- | --- |
| **Random Effects** | ID (Intercept) | 0.153 | 0.0233 | 0.390 |  |  |
|  | Residual | 0.191 | 0.0364 |  |  |  |
|  |  | **β** | **SE** | **df** | **t** | **p** |
| **Fixed Effects** | (Intercept) | 0.25 | 0.081 | 24.6 | 3.055 | **0.0054** |
|  | Treatment [GENUS] | 0.093 | 0.060 | 118. | 1.557 | 0.12 |
|  | Timepoint [30min] | -0.060 | 0.048 | 110. | -1.267 | 0.21 |
|  | Timepoint [Stim] | -0.072 | 0.059 | 116. | -1.219 | 0.23 |
|  | Days After Craniotomy | -0.0018 | 0.00070 | 33.4 | -2.609 | **0.013** |
|  | Sex [M] | -0.16 | 0.083 | 14.1 | -1.925 | 0.075 |
|  | Treatment [GENUS] x Timepoint [30min] | 0.19 | 0.080 | 110. | 2.371 | **0.019** |
|  | Treatment [GENUS] x Timepoint [Stim] | 0.23 | 0.087 | 113. | 2.642 | **0.0094** |

Supplementary Table ST7: Summary of statistical tests for Model 1: $\Delta BFI \sim Treatment \times Timepoint+Sex+(1|ID)$, with restricted maximum likelihood (REML) estimation, Satterthwaite degrees of freedom, and Tukey adjustment for multiple comparisons. Data includes 135 observations and 18 subjects (ID).

| **Comparison Type** | **Contrast** | **EMM** | **SE** | **df** | **t** | **p** |
| --- | --- | --- | --- | --- | --- | --- |
| **Effect vs. Baseline** | GENUS Stim vs. 0 | -0.012 | 0.012 | 75.6 | -0.973 | 0.33 |
|  | GENUS 0min vs. 0 | 0.020 | 0.012 | 75.6 | 1.676 | 0.098 |
|  | GENUS 30min vs. 0 | -0.00053 | 0.012 | 71.9 | -0.044 | 0.96 |
|  | Control Stim vs. 0 | -0.017 | 0.010 | 40.9 | -1.733 | 0.091 |
|  | Control 0min vs. 0 | 0.0043 | 0.010 | 40.9 | 0.438 | 0.66 |
|  | Control 30min vs. 0 | -0.0037 | 0.010 | 40.9 | -0.377 | 0.71 |
| **Effect vs. Control** | GENUS Stim vs. Control Stim | 0.0052 | 0.013 | 130 | 0.414 | 0.68 |
|  | GENUS 0min vs. Control 0min | 0.016 | 0.013 | 130 | 1.282 | 0.20 |
|  | GENUS 30min vs. Control 30min | 0.0032 | 0.012 | 130 | 0.259 | 0.80 |
| **Effect vs. Timepoint** | GENUS 0min vs. GENUS 30min | 0.021 | 0.014 | 124 | 1.531 | 0.28 |
|  | GENUS 0min vs. GENUS Stim | 0.032 | 0.014 | 124 | 2.329 | 0.056 |
|  | GENUS 30min vs. GENUS Stim | 0.011 | 0.014 | 124 | 0.828 | 0.69 |
|  | Control 0min vs. Control 30min | 0.0080 | 0.010 | 124 | 0.794 | 0.71 |
|  | Control 0min vs. Control Stim | 0.021 | 0.010 | 124 | 2.115 | 0.091 |
|  | Control 30min vs. Control Stim | 0.013 | 0.010 | 124 | 1.321 | 0.39 |

Supplementary Table ST8: Summary of statistical tests for Model 2: $\Delta VOP \sim Treatment \times Timepoint+CraniotomyAge+Sex+(1|ID)$, with restricted maximum likelihood (REML) estimation, Satterthwaite degrees of freedom, and Tukey adjustment for multiple comparisons. Data includes 135 observations and 18 subjects (ID).

| **Comparison Type** | **Contrast** | **EMM** | **SE** | **df** | **t** | **p** |
| --- | --- | --- | --- | --- | --- | --- |
| **Effect vs. Baseline** | GENUS Stim vs. 0 | 0.26 | 0.060 | 52.8 | 4.373 | **0.0001** |
|  | GENUS 0min vs. 0 | 0.10 | 0.061 | 55.6 | 1.672 | 0.10 |
|  | GENUS 30min vs. 0 | 0.23 | 0.060 | 52.8 | 3.888 | **0.0003** |
|  | Control Stim vs. 0 | -0.063 | 0.060 | 54.7 | -1.055 | 0.30 |
|  | Control 0min vs. 0 | 0.0083 | 0.050 | 31.1 | 0.167 | 0.87 |
|  | Control 30min vs. 0 | -0.052 | 0.050 | 31.1 | -1.045 | 0.30 |
| **Effect vs. Control** | GENUS Stim vs. Control Stim | 0.32 | 0.065 | 114 | 5.019 | **<.0001** |
|  | GENUS 0min vs. Control 0min | 0.093 | 0.060 | 118 | 1.557 | 0.12 |
|  | GENUS 30min vs. Control 30min | 0.28 | 0.059 | 118 | 4.835 | **<.0001** |
| **Effect vs. Timepoint** | GENUS 0min vs. GENUS 30min | -0.13 | 0.065 | 110 | -2.013 | 0.11 |
|  | GENUS 0min vs. GENUS Stim | -0.16 | 0.065 | 110 | -2.460 | **0.041** |
|  | GENUS 30min vs. GENUS Stim | -0.029 | 0.064 | 110 | -0.455 | 0.89 |
|  | Control 0min vs. Control 30min | 0.060 | 0.048 | 110 | 1.267 | 0.42 |
|  | Control 0min vs. Control Stim | 0.072 | 0.059 | 116 | 1.319 | 0.44 |
|  | Control 30min vs. Control Stim | 0.011 | 0.059 | 116 | 0.0191 | 0.98 |

Supplementary Table ST9: Summary of statistical tests for Model 3: $\Delta VOP \sim Treatment +(1|ID)$, with restricted maximum likelihood (REML) estimation, Satterthwaite degrees of freedom, and Tukey adjustment for multiple comparisons. Data includes 35 observations and 14 subjects (ID).

| **Comparison Type** | **Contrast** | **EMM** | **SE** | **df** | **t** | **p** |
| --- | --- | --- | --- | --- | --- | --- |
| **Effect vs. Baseline** | GENUS 24h vs. 0 | 0.19 | 0.091 | 27.8 | 2.128 | **0.042** |
|  | Control 24h vs. 0 | -0.055 | 0.065 | 18.1 | -0.840 | 0.41 |
| **Effect vs. Control** | GENUS 24h vs. Control 24h | -0.25 | 0.10 | 26.2 | -2.488 | **0.020** |

Supplementary Table ST10: Glossary of Allen Atlas regions identified in Figure 4.

| Acronym | Name |
| --- | --- |
| MOp | Primary motor area |
| RSPagl | Retrosplenial area, lateral agranular part |
| SSp-bfd | Primary somatosensory area, barrel field |
| SSp-un | Primary somatosensory area, unassigned |
| SSp-tr | Primary somatosensory area, trunk |
| SSp-ll | Primary somatosensory area, lower limb |
| SSp-ul | Primary somatosensory area, upper limb |
| VISpm | Posteromedial visual area |
| VISrl | Rostrolateral visual area |
| VISa | Anterior visual area |
| VISam | Anteromedial visual area |
